# CRISPR-Csx28 forms a Cas13b-activated membrane pore required for robust CRISPR-Cas adaptive immunity

**DOI:** 10.1101/2021.11.02.466367

**Authors:** Arica R. VanderWal, Jung-Un Park, Bogdan Polevoda, Elizabeth H. Kellogg, Mitchell R. O’Connell

## Abstract

Type VI CRISPR-Cas systems use the RNA-guided RNase Cas13 to defend bacteria against viruses, and some of these systems encode putative membrane proteins that have unclear roles in Cas13-mediated defense. Here we show that Csx28, of Type VI-B2 systems, forms membrane pore structures to slow cellular metabolism upon viral infection, and this activity drastically increases anti-viral defense. High-resolution cryo-EM reveals that Csx28 exists unexpectedly as a detergent-encapsulated octameric pore, and we then show these Csx28 pores are membrane localized *in vivo*. Activation of Csx28 *in vivo* strictly requires sequence-specific recognition of viral mRNAs by Cas13b, and this activation results in Csx28-mediated membrane depolarization, slowed metabolism, and inhibition of sustained viral infection. Together, our work reveals an unprecedented mechanism by which Csx28 acts as a downstream, Cas13b-activated, effector protein that uses membrane perturbation as an anti-viral defense strategy.

## Introduction

Prokaryotes use Clustered Regularly Interspaced Short Palindromic Repeats (CRISPRs) and CRISPR-associated (Cas) proteins for RNA-guided cleavage of foreign genetic elements such as viruses (bacteriophages; hence forth referred to as phage) and plasmids [1-3]. CRISPR loci possess short DNA repeats and spacer regions which bear complementarity to foreign nucleic acids [4-7]. These CRISPR loci are transcribed into CRISPR RNAs (crRNA), which complex with Cas proteins to form RNA-guided Cas complexes to detect nucleic acids of complementary sequence. Immunity is achieved by nucleic acid cleavage either through direct enzymatic degradation by these RNA-guided Cas complexes, and/or the activation of accessory nucleases [8]. Characterized into six types (I-VI), CRISPR-Cas systems are extremely diverse in target nucleic acid preference, cleavage mechanism, number of *cas* genes, and the presence of accessory genes [8]. Of interest here, Type VI systems contain a single “effector” protein, known as Cas13 (formerly C2c2), that when assembled with crRNA forms a crRNA-guided RNA-targeting complex [9, 10]. Type VI systems can be divided in four subtypes (A–D) based on the phylogeny of Cas13 [11-13], and all Cas13 proteins studied to date possess two enzymatically distinct ribonuclease activities: 1. A pre-crRNA processing nuclease that is required for mature crRNA formation, and 2. A target nuclease, comprising two HEPN domains that upon target-RNA binding form a composite active site that cleaves both foreign and host RNA transcripts indiscriminately [14]. This robust non-specific cleavage activity has been shown in several cases to lead to cellular dormancy upon targeting plasmids or phage transcripts during infection [9, 15].

Despite the identification of several novel open reading frames (i.e. accessory genes) encoded within Type VI systems, little is known about how they function [8, 11, 16, 17]. Recently, two of these accessory genes *csx27 and csx2*8 were found to significantly modulate the anti-phage defense activity of specific Cas13b- containing CRISPR systems (Type VI-B) when challenged with MS2 ssRNA phage [17], however the molecular mechanisms by which *csx27 and csx28* are able to affect Cas13b-mediated anti-phage defense are not understood. To this end, both of the putative proteins encoded by these genes were confidently predicted to contain transmembrane spanning regions [11, 17], Csx27 is potentially linked to bacterial natural competence and ubiquitin signaling systems [18], while Csx28 is predicted to contain a divergent Higher Eukaryotic and Prokaryotic Nucleotide-binding motif (HEPN) motif [11, 17], which has been hypothesized to act as an RNA nuclease [11, 14, 19, 20], however the significance of any of these predicted features is unclear. Interestingly, in a ‘guilt by association’ genetic linkage approach, it has been shown that a range of predicted membrane proteins are broadly associated with other CRISPR-Cas systems. This association suggests a more widespread, yet unexplored link, between membrane processes, CRISPR-Cas activity, and anti-phage defense [16]. More generally, membrane effector proteins have been found to exist in a range of other defense and apoptosis systems in prokaryotes, as well as more broadly across the tree of life [21-30]. However, in many cases the precise role of membrane proteins in these systems is lacking.

With this in mind, we set out to explore the potential functional association between CRISPR-Cas defense and membrane processes during phage infection. Here we focus on Cas13b and Csx28-containing Type VI-B2 systems, and we show that upon activation by target-bound Cas13b complexes during viral infection, Csx28 forms membrane pores to slow cellular metabolism, and this activity likely drastically increases anti-viral defense. Together, our work reveals an unprecedented mechanism by which a CRISPR-Cas protein uses membrane perturbation as an anti-viral defense strategy, which implies a more general link between cytoplasmic CRISPR-Cas nucleic acid detection and downstream membrane perturbation as an anti-viral defense strategy.

## Results

### Csx28 is required for optimal interference against DNA λ-phage and requires an active, phage-targeting Cas13b

We first set out to implement a phage interference system to understand at the genetic level how Csx28 contributes to Type VI-B2 anti-phage defense. As most Type VI CRISPR-Cas system spacers align to transcripts emanating from double-stranded DNA genomes (dsDNA phage or prophage) [12, 13, 17, 31, 32], we focused on the Type VI- B2 system from *Prevotella buccae ATCC 33574* (Fig. 1A) and generated synthetic CRISPR arrays containing spacers directed to the transcribed strand of early (crRNA-1), middle (crRNA-2) or late (crRNA-3) gene transcripts of the dsDNA phage lambda (λ), or a non-target control (ΔcrRNA), and subcloned these into IPTG-inducible Cas13b expression plasmids and empty plasmid controls (Fig.1B,C). These plasmids were then separately co-transformed into the laboratory *E coli* strain C3000 (which naturally lacks a Type VI-B2 CRISPR system) along with either a Csx28 expressing plasmid or an empty plasmid control (Fig. 1C). Phage susceptibility was first assessed using λ-phage Efficiency of Plating (EOP) assays. We found that Cas13b-crRNA-1 and Cas13-crRNA- 2 reduced λ-phage EOP by ∼17-fold and ∼2-fold, respectively, providing modest protection to phage infection. However, the presence of Csx28 substantially enhanced both Cas13b-crRNA-1 and Cas13b-crRNA-2 mediated anti-phage activities resulting in an additional ∼1500-fold and ∼320-fold reduction in phage infectivity (Fig. 1D).

**Fig. 1.**
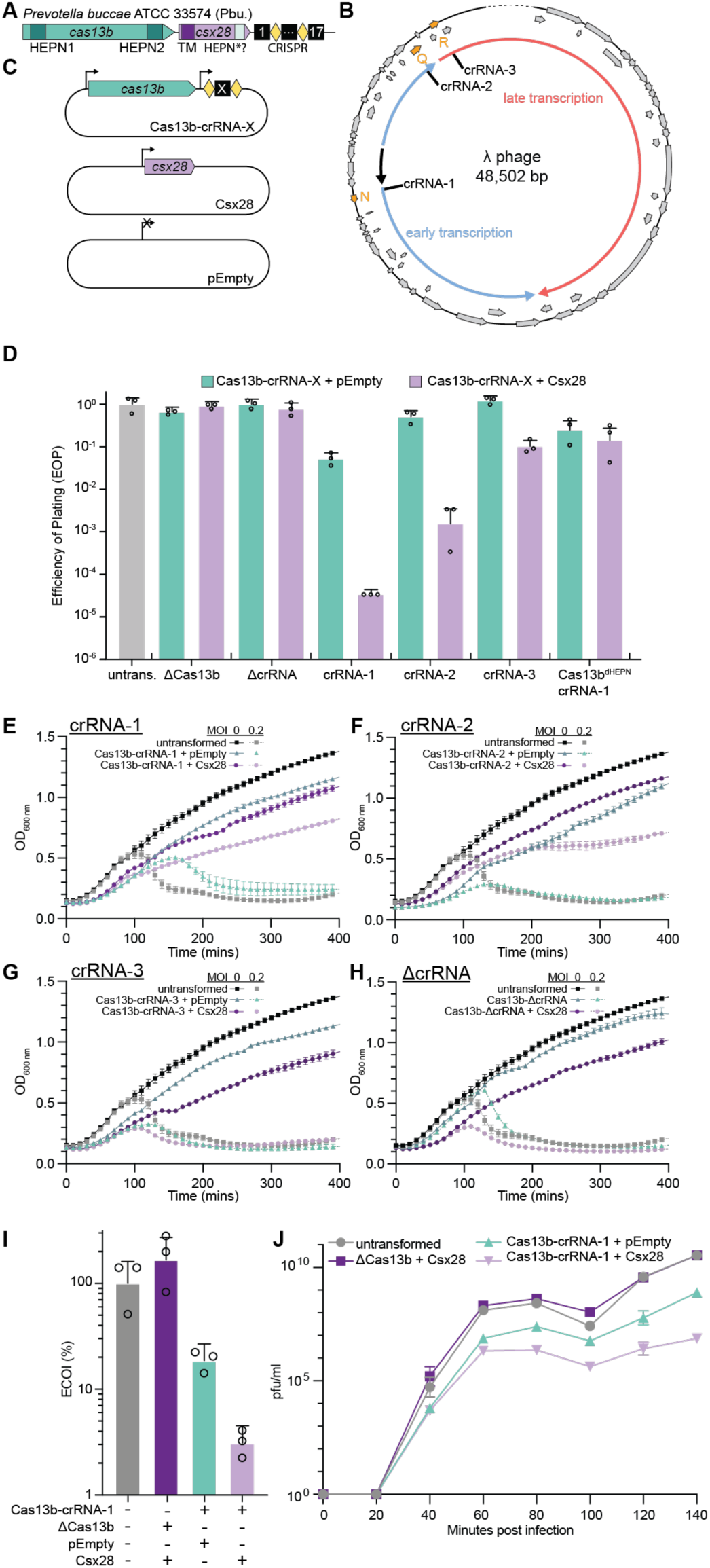
Csx28 significantly enhances Type VI-B2 CRISPR-Cas immunity against λ-phage by inducing slow-growing phenotype that helps prevent the establishment and maintenance of infection. (A) Schematic of the Type VI-B2 CRISPR-Cas system from *Prevotella buccae* used in this study, with the Cas13 and Csx28 regions of interest indicated, and the CRISPR array indicated by its repeats (yellow diamonds) and spacers (spacers rectangle). (B) Schematic of the phage λ genome in its circular form. Early and late transcription products are shown, as well as the crRNA-1 to -3 target sites. (C) Plasmid schematics for phage interference experiments in which Cas13b and Csx28 are expressed from IPTG-inducible promoters on two separate plasmids. Cas13b-crRNA-X also contains a synthetic CRISPR-Cas array driven by a constitutive promoter. pEmpty is used to control antibiotic selection pressure in these experiments. (D) Efficiency of Plating (EOP) assays measuring λ-phage infection susceptibility of untransformed (untrans.) *E. coli* or strains carrying the indicated Cas13b-crRNA-X, Csx28, or pEmpty plasmids. ΔcrRNA is a non-targeting crRNA control, ΔCas13b is no Cas13b control, Cas13b^dHEPN^ is a HEPN-nuclease deactivated Cas13b control. (E-H) Growth curves of *E. coli* strains carrying the indicated Cas13b-crRNA-X, Csx28, or pEmpty plasmids, as measured using OD600 after the addition of λ-phage at an MOI of 0.2. (E-H) are growth curves for crRNA-1, crRNA-2, crRNA-3 and ΔcrRNA- containing strains, respectively. (I) Efficiency of Center of Infection (ECOI) assays measuring λ-phage infective center formation of *E. coli* strains carrying the indicated Cas13b-crRNA-X, Csx28-containing, or pEmpty plasmids infected with λ-phage at an MOI of 0.1. (J) Phage growth (burst) assays measuring the λ-phage production over time for *E. coli* strains carrying the indicated Cas13b-crRNA-1, Csx28-containing, or pEmpty plasmids infected with λ-phage at an MOI of 0.1. Data in (D-J) are shown as mean ± s.e.m for *n* = 3 biological replicates.

Importantly, this robust Csx28-mediated enhancement of anti-phage defense requires the presence of an active, λ -targeting Cas13. The absence of Cas13 (ΔCas13), the absence of a λ -targeting crRNA (ΔcrRNA), or the mutation of Cas13’s HEPN nuclease active site residues (Cas13b^dHEPN^) completely abrogate this Csx28-mediated anti-phage enhancement effect (Fig. 1D). These results recapitulate a similar Csx28 anti-phage defense effect observed in MS2 ssRNA phage experiments [17]. We additionally observed that enhanced anti-MS2 phage defense absolutely requires a targeting Cas13b, as Csx28 expressed alone doesn’t offer any measurable defense (Fig. S1).

To obtain a more detailed insight into the kinetics of Csx28-enhanced λ-phage interference and its effect on bacterial growth at the early stages of infection, we monitored bacterial growth rates post phage infection (Fig. 1E-H). Much like in our EOP assays, Cas13b-crRNA-1 and -crRNA-2 can respond to λ-phage infection at a multiplicity of infection (MOI) of 0.2 and result in delayed lysis at the population level, cessation of growth, and a reduction but not complete loss of cell numbers (Fig. 1E, F). In contrast, crRNA-3 and ΔcrRNA respond similarly to untransformed *E. coli* (Fig. 1G, H). In the case of Cas13b-crRNA-1 and -crRNA-2, the addition of Csx28 can rescue this defect resulting in continued, albeit slower growth relative to infected cells. This suggests that Csx28 is acting to prevent phage propagation and/or cell lysis thereby enabling the cultures to continue to increase in cell numbers (Fig. 1E, F). Unsurprisingly, while the Csx28 enhancement effect is significantly muted at a higher MOIs (MOI of 2), Cas13b-crRNA-1 in the presence of Csx28 still can resist cell death post λ-phage infection (Fig. S2).

To determine at what stage of λ-phage infection Csx28 is acting to enhance defense, we carried out efficiency of center of infection (ECOI) assays and phage growth assays. The ECOI assays, which measure the percentage of initially infected cells that go on to release at least one newly synthesized phage progeny, revealed that Cas13b:crRNA-1 expression alone resulted in ∼18.5% of infected cells releasing at least one infectious virion, and the addition of Csx28 to Cas13b:crRNA-1 cultures (but not Csx28 strains alone) further reduced the release of phage to only ∼3% of infected cells. This indicates that Csx28 can enhance defense by limiting the number of initially infected cells releasing phage progeny. (Fig. 1I). To observe phage accumulation within our system, the first viral burst size was determined for each host and showed a general reduction of phage numbers per infected cell when hosts were protected with Cas13b and Csx28 compared to untransformed *E. coli* (Fig. S3). Reduction of functional phage virions within bacteria containing Cas13b:crRNA-1 and Csx28, was further amplified in subsequent time points with ∼20 and ∼140 fold average decreases in phage numbers released per infected cell compared to Cas13b only and untransformed *E. coli*, respectively, during the 2nd burst (140 minutes post infection) (Fig. 1J). Taken together, it is clear that an actively targeting Cas13b is required to activate Csx28 which allows for robust anti-phage defense response against a dsDNA phage, and this is possibly achieved by Csx28 inducing a slow growing phenotype that helps prevent the establishment and maintenance of λ-phage infection.

### Cryo-EM reveals that Csx28 forms an octameric membrane pore

To further understand how Csx28 is functioning to enhance Cas13b-mediated antiphage defense, we expressed and purified recombinant Csx28 from *E coli*. Despite previous evidence that indicated Csx28 is uniformly expressed in the cytosol [17], we found that Csx28 was completely insoluble in standard cytosolic protein purification buffers and required detergent solubilization, suggesting it may in fact be membrane-associated *in vivo*. Out of a panel of commonly used membrane protein detergents, we found that dodecyl maltoside (DDM) resulted in the highest yields and purity of soluble Csx28. Size exclusion chromatography suggested DDM-solubilized Csx28 was forming several discrete, non-exchanging oligomers of different sizes in solution (Fig. S4; henceforth referred to as light and heavy fractions). To precisely measure the molecular weight of these oligomers, we turned to static-light scattering coupled with size-exclusion chromatography (SEC-SLS). We observed that the heavy fraction of Csx28 forms discrete octamers of ∼170 kDa (monomeric Csx28 has a molecular weight of ∼21 kDa), that were dynamically exchanging with a larger 16-mer species during the experiment (Fig. 2A, Fig. S5). Unfortunately, because the light fraction coelutes with empty DDM micelles, calculating an accurate experimental molecular weight of this Csx28 complex wasn’t possible using this technique.

**Fig 2.**
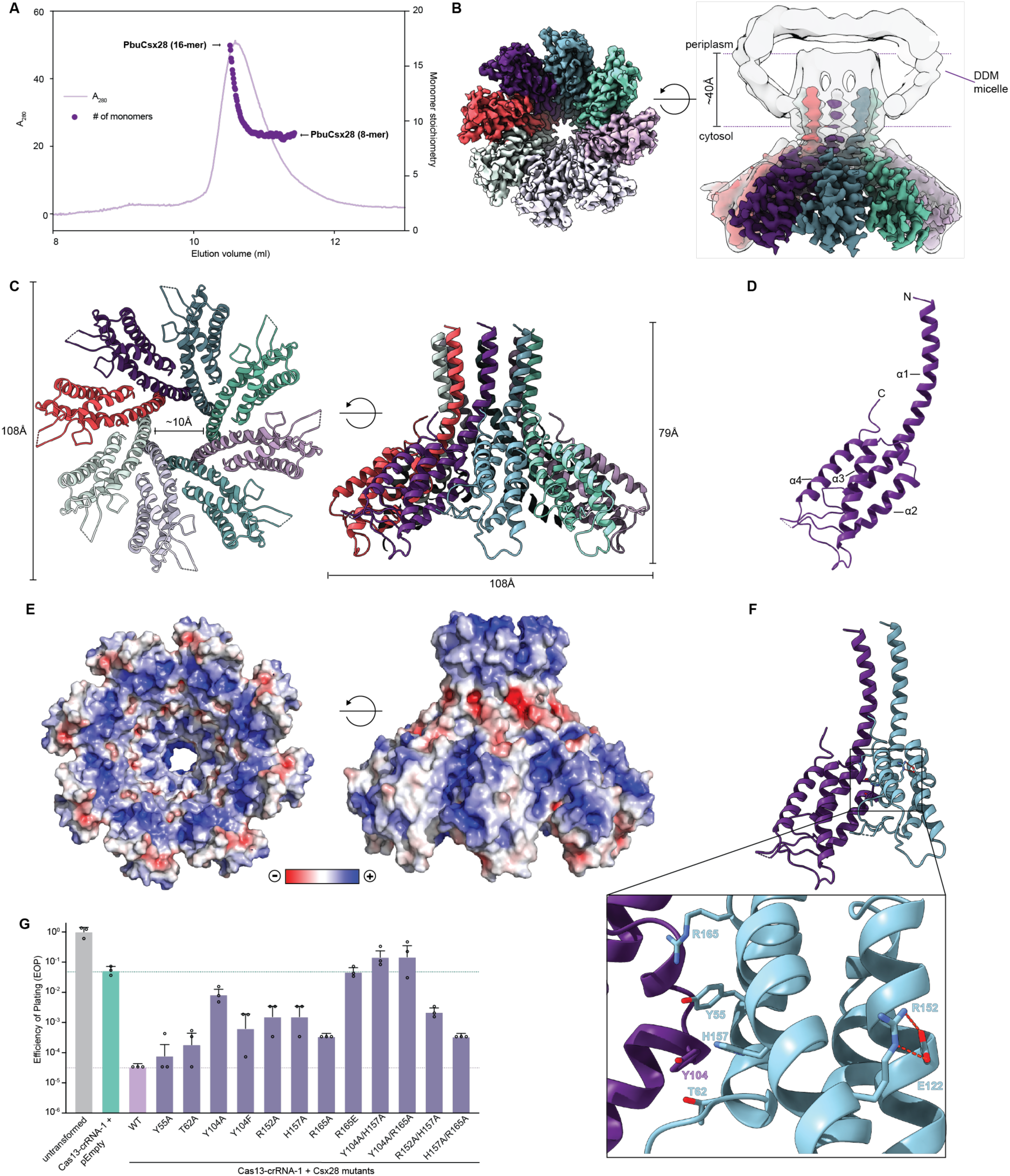
Cryo-EM reveals that Csx28 forms an octameric membrane pore with a unique protomer interface. **(A)** Static light scattering coupled with size-exclusion chromatography analysis of Csx28 heavy fraction. Absorbance at 280nm (A280) is plotted as a purple line and Csx28 monomer stoichiometry across the Csx28 peak is plotted as purple circles. Please see fig. S5 for full three detector traces of Csx28 and a BSA standard. Please note that under these experimental conditions Csx28 undergoes dynamic exchange between a 16-mer and 8-mer, which results in a weighted average oligomeric state across the peak. (**B**) High resolution (3.65Å) cryo-EM reconstruction of Csx28 (with each identical protomer colored uniquely) embedded in a DDM micelle, which is displayed as a composite high-resolution cryo-EM map superimposed with a 8 Å low-pass filtered version of the same map to display lower resolution features, such as the DDM micelle and transmembrane helices. The predicted TMHMM orientation of Csx28 within in the cytoplasmic membrane and the predicted width of the membrane is shown for reference. (**C**) Bottom and side views of the atomic model of the Csx28 octamer with each identical protomer uniquely colored. The dimensions of the octamer and the diameter at the constriction of the pore are also shown. **(D)** Atomic model of an isolated Csx28 protomer with each helix of the four alpha-helical bundle labeled. **(E)** Electrostatic surface representations of the bottom and side of Csx28. The red-blue color gradient represents negative to positive electrostatic potential (± 5kT/e). (**F**) A magnified view of the Csx28 protomer-protomer interface. Amino acid side chains that form the interface with the divergent HEPN motif residue H157 are shown as sticks and labeled. Divergent HEPN motif residue R165 is shown forming a salt bridge with E122. (**G**) Efficiency of Plating (EOP) assays measuring the effect of amino acid mutations at the Csx28 protomer-promoter interface on λ-phage infection susceptibility of *E. coli* strains carrying the indicated Cas13b-crRNA-1 and Csx28 (WT or mutant) plasmids. Data is shown as mean ± s.e.m for *n* = 3 biological replicates.

Given this observation, we decided to pursue cryo-EM structural studies and were able to determine the structure of Csx28 (heavy fraction) embedded in a DDM micelle to an estimated global resolution of 3.65Å (Fig. 2B, Fig. S6, Table S1.). 2D class averages clearly indicated the presence of 8-fold symmetry; imposition of C8 symmetry resulted in the high-resolution cryo-EM reconstruction shown (Figure 2B). The resulting reconstruction is a homo-octamer with an 8-fold symmetry about a central pore; a nearly full-length model, corresponding to residues 19-171, was built into the asymmetric unit (full length Csx28 is 177 amino acids). The structure can be divided into two distinct regions: a partially unresolved single N-terminal alpha helix that is embedded within a DDM micelle (matching the membrane topology prediction generated by TMHMM [33]), and a well-ordered C-terminal cytoplasmic domain (Fig. 2B,C). As commonly observed in membrane protein structures, the DDM micelle appears as diffuse, spherical density (Fig. 2B), the remaining low-resolution features apparent in a low-pass filtered version of the cryo-EM map indicates how Csx28’s N-terminal transmembrane helix may traverse the lipid bilayer. Interestingly, the 3D class averages perfectly recapitulate our SEC-SLS data, with the two major classes forming octamers and the minor class forming a 16-mer (Fig. S6). The central pore has a minimal diameter of approximately 10Å, a similar to the diameters observed in many large-pore forming channels (e.g. connexin gap junction channels), that permeate ions and small metabolites such as ATP but is likely too small for the passage of proteins such as phage endolysins [34]. Each protomer is organized as a four-helix bundle (ɑ1-4), with the N-terminal helices (ɑ1-2) lining the inside of the pore with the two C-terminal helices (ɑ1-2) forming the outside of the pore (Fig. 2D). Neither a DALI search [35] or Omakage [36] search found any deposited structures with significant structural similarity to a single Csx28 monomer, or overall shape similarity to the Csx28 octamer, respectively.

The Csx28 protomers are arranged in a parallel head-to-head orientation (Fig. 2C) and this arrangement results in a pore lined with mostly positively charged amino acid side chains, as visualized by electrostatic surface potential maps (Fig. 2E). These positively charged regions may provide selectivity towards specific ions or metabolites, or act as potential nucleic acid binding sites, especially given Csx28 was previously predicted to contain a divergent HEPN motif, which may act as RNA-binding and/or cleavage domain [17]. Canonical HEPN motif containing proteins often form “face-to-face” dimers that result in the HEPN motif from each protomer facing towards each other, lining the dimer interface. This dimer interface often acts as an RNA-binding surface and/or RNase active site [27]. While Csx28 adopts a similar four alpha-helical bundle fold, common to previously determined HEPN motif containing proteins, the oligomers form a noncanonical “face-to-back” arrangement, which results in only one HEPN motif per interface, rather than the expected two observed in canonical HEPN-containing proteins. In this Csx28 structure, only one of the predicted HEPN motif (RX_4-6_H) residues, H157, (Fig. 2F) and a distinct set of conserved residues from a neighboring alpha helix and protomer (e.g. Y55, Y104, T62, R165) form the interface. In addition, the predicted HEPN-motif arginine (R152), canonically required for RNA hydrolysis, is oriented 180° away from the interface and is engaged in a salt bridge with E122 from helix ɑ2 in the same protomer.

To further probe whether the protomer-protomer interface was required for phage defense activity, we generated several single- and double-point mutations within this interface. We found that single point mutations in this region resulted in a ∼2- to ∼1400- fold reduction in the ability for Csx28 to enhance λ-phage defense, and double point mutations were able in most cases to further exacerbate this effect (Fig. 2G).These results suggest that small changes within the interface region can significantly affect Csx28’s ability to function in phage defense. We also probed the importance of the R152:E122 salt bridge, that we observe sequestering the predicted HEPN arginine away from the interface, by performing a charge swap analysis. We observed that single point mutants R152E and E122R result in a loss of Csx28-mediated λ-phage defense. However, combining these two mutations, with the idea of rescuing salt bridge formation, results in almost complete rescue in λ-phage defense (Fig. S7) which indicates that this salt bridge is important for maintaining the structure of the Csx28 protomer and maintaining the structurally integrity of the complex.

### Csx28 forms membrane-localized oligomers in vivo that result in significant membrane depolarization and reduced metabolism upon Cas13b activation

Given the detergent embedded, pore-like structure of Csx28 we observed in our cryo-EM analysis, we questioned how Csx28 may be affecting membrane function *in vivo*. To start, we first wanted to observe the cellular localization of Csx28 and Cas13b expressed in *E. coli*. To do this experiment, we tested a range of small-epitope tagged Csx28 and Cas13b constructs using EOP assays to ensure that the addition of a tag did not affect Csx28 and Cas13’s function. Importantly, we found that while the addition of a C-terminal V5 tag or an N-terminal StrepII tag did not impact Csx28 activity, we saw a loss of activity when tagging with the highly charged FLAG epitope tag (Fig. S8A), highlighting the importance of appropriate tag selection for this protein. Similarly, we saw differences in activity depending on which terminus on Cas13b we added a 3xHA tag (Fig. S8B), and we decided that the combination of C-terminal V5 tag for Csx28 and an N-terminal 3xHA tag for Cas13b was the best choice moving forward.

With these tagged proteins in hand, we used an ultra-centrifugation and DDM detergent solubilization based membrane fractionation strategy coupled with Western blotting to determine localization of Cas13b and Csx28. Cas13b was found to co-fractionate with DnaK (a cytosolic chaperone) in the cytosol, whereas Csx28 was found to exclusively reside in the DDM-soluble membrane fraction, co-fractionating with OmpC (a membrane porin) (Fig. 3A). Then, we sought to determine whether Csx28 can form oligomers *in vivo*. Cas13b and Csx28- expressing *E. coli* were treated with a membrane permeable protein crosslinker disuccinimidyl suberate (DSS) pre- and post- λ-phage infection followed by Western blot analysis on total cell lysates. We observe that indeed Csx28 exists in a potential ensemble of oligomers in growing *E. coli* (Fig. S9), with a protein molecular weight banding pattern that closely resembles the crosslinking of increasing multiples of Csx28 monomers. This result emphasizes that the octameric form we observe in our cryo-EM structure can likely form in native *E coli* membranes.

**Fig. 3.**
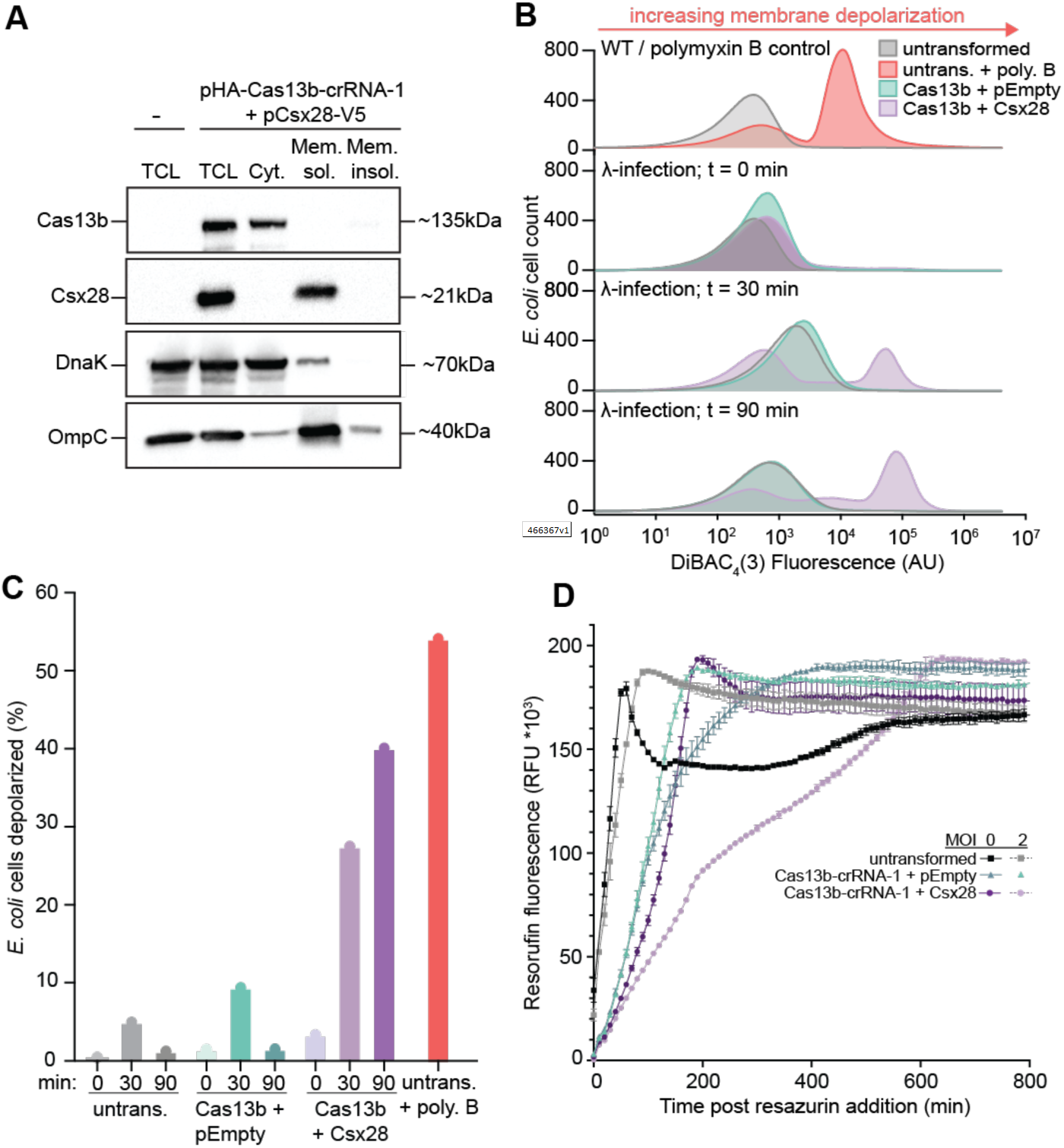
Csx28 localizes to the membrane in E. coli, acts to cause membrane depolarization and a loss of metabolic activity upon Cas13b activation during λ-phage infection. (**A**) Western blot to detect the localization of Cas13 and Csx28 cytosolic vs. detergent-soluble and detergent-insoluble fractions obtained from *E. coli* expressing HA-tagged Cas13b and/or V5-tagged Csx28. Blots were first probed with anti-HA or anti-V5 antibodies to detect HA-Cas13 and Csx28-V5, respectively. Blots were also probed for DnaK and OmpC as cytosolic and membrane fractionation controls, respectively. (**B**) Flow cytometry histograms of a DiBAC4(3) staining assay measuring membrane depolarization of WT *E. coli or E. coli* strains expressing Cas13b-crRNA-1 or Cas13b-crRNA-1 and Csx28 over the course of a λ-phage infection (MOI of 1). An increase in DiBAC4(3) fluorescence represents an increase in membrane depolarization. A Polymyxin B (Poly. B; a membrane integrity disrupter) treated *E. coli* sample was used as positive control for membrane depolarization. In all samples, 100,000 *E. coli* cells were analyzed. (**C**) Quantification of the percentage of depolarized cells in (C), determined by calculating the area under the curve of the depolarized cell subpopulation (i.e., number of cells depolarized) as a percentage of the total population. (**D**) Resazurin metabolism assay measuring the conversion of resazurin to fluorescent resorufin over time as a function of active respiration for untransformed *E. coli* or strains expressing Cas13b-crRNA-1 or Cas13b-crRNA-1 and Csx28 either in the absence or presence of λ-phage infection (MOI of 2). Resazurin is added to the growing cultures at one hour post λ-phage infection. Data is shown as mean ± s.e.m for *n* = 3 biological replicates.

The combined observation of an octameric Csx28 pore by cryo-EM, membrane-localized Csx28 oligomer formation *in vivo* and a slow growing Cas13b:crRNA1-Csx28 phenotype during phage infection, led us to wonder whether the oligomerization of Csx28 *in vivo* results in membrane pores that may perturb the basal electrochemical potential that exists across *E. coli* cell membranes, knowing that this potential is an essential requirement for aerobic metabolism. To test this hypothesis, we employed a membrane depolarization assay that uses the cationic dye Bis-(1,3-Dibutylbarbituric Acid) Trimethine Oxonol (DiBAC_4_(3)), which only becomes fluorescent after accumulating in cells that have de-energized membranes due to a loss of membrane potential [37]. To carry out this experiment, Cas13b:crRNA-1/Csx28-containing strains were first infected with λ-phage and culture samples were collected and treated with DiBAC_4_(3) at 0, 30, and 90 minutes post infection. DiBAC_4_(3) fluorescence was then measured using flow cytometry to ascertain membrane polarization on a per cell basis. Strikingly, only Cas13b:crRNA-1/Csx28 resulted in pronounced membrane depolarization with as much as 40% of the population depolarized at 90 minutes post infection (Fig. 3B,C), while expression of Cas13b:crRNA-1 (Fig. 3B), Csx28, or Cas13b:ΔcrRNA (Fig. S10A) alone did not result in dramatic increases in membrane depolarization (Fig. 3B-C). As a control, treating untransformed *E. coli* with polymyxin B, an antibiotic that is known to disrupt membrane integrity, resulted in a similar depolarization phenomenon as Cas13b:crRNA-1/Csx28 (Fig 3B-C). To investigate whether this Csx28-mediated depolarization resulted in larger defects in membrane integrity, we performed the same flow cytometry experiments but stained the cells with membrane-impermeable DNA-stain propidium iodide (PI), a dye that requires more gross defects in membrane integrity to observe positive staining. We observed that Csx28 activation did not result in large changes in PI fluorescence, relative to a membrane disrupting antibiotic polymyxin B (Fig. S10B), suggesting that whatever pores Csx28 may be forming likely do not result in drastic loss of the structural integrity of the membrane. Taken together, our depolarization observations are in line with the slow-growing phenotype observed in figures 1E-F and figure S2, and with previous studies showing that *E. coli* can survive and continue to growth post transient membrane depolarization [38, 39].

Finally, we carried out resazurin assays to test whether Csx28 elicited membrane depolarization affects cellular metabolism. Resazurin is a non-fluorescent substrate that is irreversibly converted by NADH/NAPDH dependent dehydrogenases to the fluorescent product resorufin only in actively respiring cells that have sufficient NADH/NAPDH pools, thus can be used as a measure of cellular respiration rates [40]. Untransformed *E*.*coli*, Cas13b:crRNA-1, Csx28, and Cas13b:crRNA-1/Csx28- containing strains were infected with λ-phage (MOI of 2, Fig. 3D; MOI of 0.2, Fig. S11) for 60 minutes (to complete approximately one infection cycle) before resazurin was added to the cultures and fluorescence measurements were collected over 800 mins. We observed that most of the cultures were able to completely metabolize resazurin to resorufin within ∼300 minutes, even in the presence of a phage infection and the subsequent crash of the cell population. However, cultures containing Cas13b:crRNA- 1/Csx28 exhibited markedly different resazurin turnover kinetics, with two phases of noticeably slower turnover, which resulted in these cultures requiring ∼600 minutes to completely turnover resazurin. This much slower rate of resazurin turnover is highly indicative of a reduced rate of metabolism, likely caused by the loss of membrane polarization induced by activated Csx28 pore formation. We hypothesize that attenuated metabolic rate allows the cultures to access a cellular state that overcomes chronic phage infection. Collectively, these results indicate that *in vivo*, Csx28 membrane oligomers are activated by targeting Cas13b which results in Csx28 pore formation and subsequent depolarization of the cytoplasmic membrane, which leads a cellular state marked by a distinctly slow metabolic rate and slowed culture growth.

## Discussion

Here we have shown that Csx28 forms an octameric membrane pore that upon activation by Cas13b is able to substantially enhance the ability of Type VI-B2 CRISPR-Cas systems to mount an immune response against both dsDNA and RNA phage. We show genetically that Csx28 activation and subsequent anti-phage defense requires an active targeting Cas13b:crRNA complex, and that inactivating the HEPN nuclease domain of Cas13 completely abrogates the activation of Csx28-enhanced anti-phage defense. We also show that Csx28 activation upon Cas13b:crRNA-sensed phage infection results in a slow growing phenotype, as a result of Csx28 pore formation, concomitant membrane depolarization, and slowed cellular metabolism.

Based on these data, we propose the following model for Csx28 activation and activity (Fig. 4): Csx28 is recruited to the membrane and exists in an inactive state comprising possibly a dynamic ensemble of Csx28 multimers, whereas Cas13b:crRNA complexes reside in the cytoplasm surveying any RNA transcripts that may bear complimentary to loaded crRNAs. Upon phage infection and successful detection of phage transcripts by Cas13:crRNA complexes, Cas13 undergoes a conformational change that activates it’s HEPN nuclease for collateral ssRNA cleavage, which results in non-specific RNA degradation of both viral and possibly host transcripts. We speculate that the subsequent activation of Csx28 and the formation of an open membrane pore and membrane depolarization occurs via either: (a) Csx28 is activated via an interaction with Cas13-generated RNA cleavage products, which possess noncanonical RNA termini that may act as a specific signal for Csx28 activation. Activation of Csx28 via Cas13b cleavage products would be similar to what is observed with cyclic oligoadenylate (cOA) signaling within Type III CRISPR-Cas systems [41], or damaged tRNA halves as specific signaling activators of RNA repair systems [42]; or (b) a direct binding interaction between Csx28 and Cas13:crRNA:target-RNA that can only be formed when Cas13 is in this ternary conformation. This protein-protein interaction between Csx28 and activated Cas13 could be similar to what is observed within Type I CRISPR-Cas systems when the target-bound Cascade complex recruits the effector DNase, Cas3 [43-45].

**Fig. 4.**
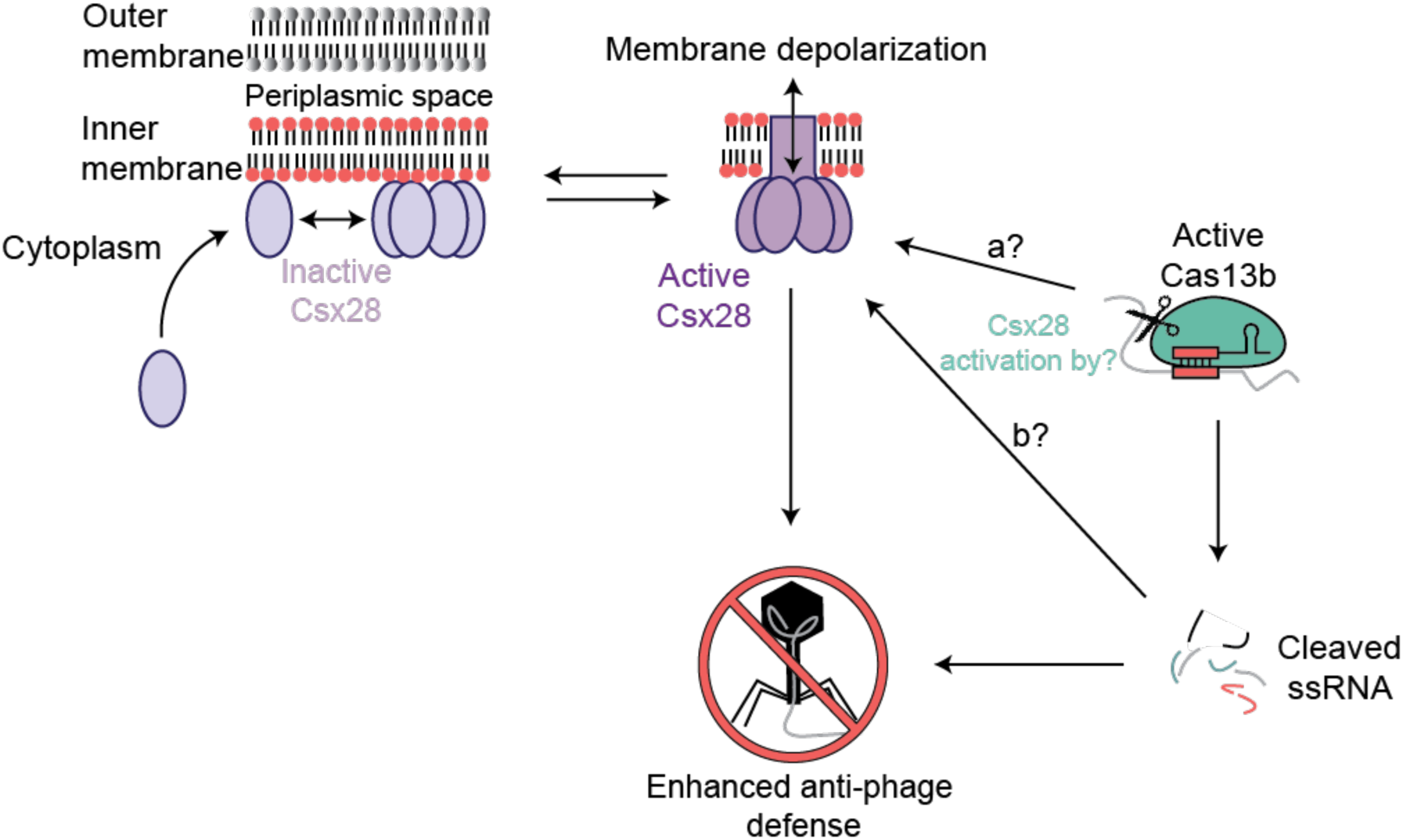
A mechanic model describing Csx28’s role in enhancing TypeVI-B2 anti-phage defense. Csx28 is recruited to the membrane prior to phage infection and exists in an inactive state, whereas Cas13b:crRNA complexes reside in the cytoplasm. Upon successful detection of phage transcripts by Cas13:crRNA complexes, Cas13’s HEPN nuclease is activated for collateral ssRNA cleavage, which results in non-specific RNA degradation of both viral and host transcripts. The subsequent activation of Csx28 and the formation of an open membrane pore and subsequent membrane depolarization occurs likely via either: (a) a direct binding interaction between Csx28 and HEPN-activated Cas13:crRNA:target-RNA or (b) Csx28 is activated via an interaction with Cas13-generated RNA cleavage products, which possess non canonical RNA termini that may act as a specific signal for Csx28 activation.

Structurally, Csx28 represents a new class of pore-forming membrane protein, as it possesses no noticeable single-chain structural or overall shape similarity to any previously determined protein structures. We hypothesize the oligomeric Csx28 structure we capture in a DDM micelle represents an open, membrane permeable pore. Furthermore, while Csx28 was hypothesized to possess a divergent HEPN RNA-binding and/or RNase motif [11, 14, 17, 19, 20], the ‘face-to-back’ interface formed between the Csx28 protomers in our structure renders any potential HEPN interface in an inactive nuclease form. The inactive form observed here is somewhat reminiscent of what is observed with RnlA, a HEPN-nuclease containing toxin in it’s in resting, nuclease- inactive state [46]. However, unlike RnlA which can switch between a canonical active face-to-face HEPN interface and a resting ‘back-to-back’ interface, from our mutagenesis data, we hypothesize that the divergent ‘face-to-back’ interface formed by Csx28 is the active state required for anti-phage defense. Specifically, in our current model, we hypothesize that divergent interface is required for the formation of a functional, active membrane pore, and the structural integrity of this octameric pore is required for phage defense. However, based on our model, one could envisage that RNA binding within this region may seed the formation of active Csx28 pores. Relatedly, with respect to pore formation, our structure of Csx28 also provides strong evidence that the N-terminal helix forms a functional transmembrane spanning region, the same region as correctly predicted by the membrane topology algorithm TMHMM [33].

Membrane pore proteins appear to be a common feature of prokaryotic immune systems, as well as various defense and apoptosis-inducing systems across the tree of life. For example, many described prokaryotic systems including phage exclusion systems such as the Rex system [47], a subset of TA, Thoreis, Zorya, Kiwa, CBASS and Retron defense systems, as well as hypothetical systems that contain nucleotide- synthesizing enzymes, have been found to contain potential pore-forming effectors [21-30]. In addition, a range of potential pore-forming effectors, e.g. the cation transporter CorA, and a number of hypothetical membrane-associated CARF-domain containing proteins are also found embedded within a number of CRISPR-Cas systems (especially Type III CRISPR-Cas systems). It has been hypothesized these predicted membrane proteins are involved in the regulation CRISPR-Cas defense [16]. Our current data suggests that rather than acting to stimulate the RNase activity of the associated Cas13b as previously hypothesized [17], Csx28 conversely acts as a terminal effector that is activated by Cas13b in anti-phage defense, which presents a possible mechanism for other hypothetical CRISPR-Cas and defense-linked membrane proteins. It is tantalizing to speculate that the RNA-guided nucleases in many of these defense systems, have been co-opted as nucleic-acid sensors that relay the presence of foreign nucleic acid to terminal membrane protein effectors. This signaling process allows for a rapid promotion of an abortive infection phenotype, which could be either a slowed growth cellular metabolic state or cell death, neither of which are conducive to optimal phage replication. Taken together, our work provides a conceptual starting point to understand how RNA-guided CRISPR-Cas nucleases can potentially act more broadly as nucleic-acid sensors that signal infection status to downstream membrane effector proteins, which in turn perturb membrane integrity as an anti-viral defense strategy, highlighting a potentially general and underappreciated aspect of CRISPR-Cas systems.

## Materials and Methods

### Bacterial Strains and Phages

*E.coli* C3000 (ATCC 15597) were grown in LB or 1.5% LB agar at 37 °C shaking at 200 rpm. Whenever applicable, media were supplemented with ampicillin (50 μg ml^−1^) and/or chloramphenicol (15 μg ml^−1^) to ensure the maintenance of plasmids. C3000 cells were used to propagate MS2 phage (ATCC 15597-B1) according to ATCC recommended handling procedure.

*E.coli* MG1655 harboring phage λ cI857 bor::kanR (a heat inducible mutant prophage; [48]) were grown in LB supplemented with 10 mM MgCl_2_ and kanamycin (50 µg ml^-1^) at 30 °C shaking at 200rpm until an OD of 0.4. The temperature was increased to 42 °C to elicit the lytic response of the phage which required ∼2 hours for complete clearance of the medium. λ-phage was then purified using a previously published protocol [49] and stored in SM buffer (50 mM Tris-Cl pH 7.5, 100 mM NaCl, 8 mM MgSO_4_) at 4 °C. *E.coli* Rosetta2 (DE3) pLysS (Novagen) transformants were grown in terrific broth (TB) at 37 °C shaking at 180rpm with ampicillin (50 μg ml^−1^) and chloramphenicol (15 μg ml^−1^) to ensure the maintenance of plasmids.

### Plasmid Construction

Csx28 and Cas13b plasmids in this study are modified versions of the following Addgene plasmids #89909 and #89906, respectively. These plasmids were modified to include a lac operator sequence which is necessary for functional lac operon induction in the presence of IPTG and suppression in the presence of glucose. Additionally, crRNA spacer sequences were inserted between direct repeats at the BsaI restriction site of Cas13b plasmids. Mutants of Csx28 and Cas13b were generated via site-directed mutagenesis.

### Efficiency of Plating (EOP)

λ phage was titrated by ten-fold serial dilutions in phage SM buffer. 2 µl of each serial dilution were spotted onto solidified 0.75% top LBA containing 200 µl *E. coli* culture (pre-grown in 5 ml LB overnight) in 3ml top LBA. Plates were incubated at 37 °C for 16 hours. Plaques were counted and titer was determined as plaque forming units (pfu) ml^−1^. EOP was calculated as (pfu ml^−1^ (test strain)/pfu ml^−1^ (control strain, *E. coli* C3000)). All experiments include three individual biological repeats with each including technical replicates in triplicate. Csx28 and Cas13b and plasmid mutants were transformed into *E. coli* C3000 prior to plating. Data was plotted using GraphPad Prism 9.

### One-step phage growth curves

Overnight solutions of *E. coli* were used to inoculate 20 ml cultures of LB+10mM MgCl_2_ at a starting OD_600_ of 0.1 and were grown to OD_600_ of 0.4 when they were infected with λ phage at an MOI of 0.1. Adsorption was carried out for 5 min, followed by three 20ml washes with LB+10mM MgCl_2_ to remove “free” unabsorbed phage. Pellets were resuspended in 20 ml LB+10mM MgCl_2_. For the 0min time point (Efficiency of Center of Infection, ECOI), two 100µl aliquots were taken immediately: one serially diluted in LB+10 mM MgCl_2_ (untreated sample) and plated directly on top LBA with susceptible host *E.coli*, and the other 100 µl aliquot was treated with a 2% chloroform LB+10mM MgCl_2_ solution (treated sample; chloroform lyses the cells and results in only the measurement of mature phage) for 2-3 mins and then serially diluted in LB+10 mM MgCl_2_ and plated on top LBA with *E.coli*. Subsequent time points were subjected to only chloroform treatment and titrated in the same manner. At each time point, the pfu ml^−1^ was determined for each strain with the titer representing the number of infectious centers formed. The ECOI was calculated from the untreated sample at 0 min as (pfu ml^−1^ (test strain)/pfu ml^−1^ (control strain, *E. coli*)). The average first burst size was calculated as (pfu ml^-1^ (80min treated)/ECOI). All experiments include three individual biological repeats with each including technical replicates in triplicate. Data was plotted using GraphPad Prism 9.

### Bacterial Growth Curves

Overnight cultures of bacteria (*E. coli* with plasmids pEmpty, Cas13b, and/or Csx28) or negative control (*E. coli* with both empty plasmids) were diluted to a final OD_600_ 0.1 in LB+10 mM MgCl_2_ supplemented with ampicillin and chloramphenicol in presence of λ phage for a final MOI of 0, 0.2 and 2. 200 μl of the mixed samples were transferred into wells of a 96-well plate in triplicate. Plates were incubated at 37 °C with shaking in a Tecan Spark plate reader with A_600_ measurements taken every 10 minutes with the first time point at 0 min. Data was plotted using GraphPad Prism 9.

### Csx28 protein expression and purification

The Csx28 expression vector was assembled by using a PCR fragment of the PbuCsx28 ORF from the PbuCsx28 Addgene plasmid (#89909; Table S2). Csx28 was C-terminally tagged with a TEV-MBP-His6 cleavage site sequence with expression driven by a T7 promoter. Expression vectors were transformed into Rosetta2 (DE3) pLysS *E. coli* cells and grown in TB broth at 37 °C, induced at mid-log phase with 0.5 mM IPTG, and grown for an additional three hours until harvested. Cell pellets were resuspended in lysis buffer (50 mM HEPES pH 7.5, 1 M NaCl, 5% glycerol, 1 mM TCEP, 1% DDM, 20mM imidazole, and EDTA-free protease inhibitor (Roche)), lysed by sonication, and clarified by centrifugation at 16,000g. Soluble Csx28-TEV-MBP-His6 was isolated over metal ion affinity chromatography and rinsed with wash buffer (50mM HEPES pH 7.5, 1 M NaCl, 5% glycerol, 1 mM TCEP, 0.1% DDM, and 40mM imidazole) and eluted in elution buffer (50mM HEPES pH 7.5, 1 M NaCl, 5% glycerol, 1 mM TCEP, 0.1% DDM, and 300mM imidazole). Eluate was treated with TEV protease at 4 °C during overnight dialysis in dialysis buffer (50mM HEPES pH 7.5, 500 mM NaCl, 5% glycerol, and 1 mM TCEP) to remove the MBP-His6 tag. Cleaved protein was loaded onto a second metal ion affinity column to clear the MBP-His6 tag. Csx28 eluate was concentrated and further purified via size-exclusion chromatography on a GE S200 column in gel filtration buffer (20mM HEPES pH 7.5, 200 mM KCl, 1 mM TCEP, 5% glycerol, and 0.1% DDM) and stored at −80 °C. Size exclusion chromatography traces were plotted using GraphPad Prism 9.

### Static light scattering size exclusion chromatography (SEC-SLS)

Purified Csx28 was loaded onto a Superdex 200 increase column (GE Healthcare) equipped with in-line UV (Postnova Analytics), static light scattering and refractive index detectors (Precision Detectors), and equilibrated with a buffer containing 20mM HEPES pH 7.5, 200 mM KCl, 1 mM TCEP, 5% glycerol, and 0.1% DDM. Data were collected using (PrecisionAcquire32 and PrecisionDiscovery32, Precision Detectors), and molecular masses were calculated using manually normalized values and the three detector method [50] using BSA as a known molecular mass control. Data was plotted using GraphPad Prism 9.

### Cryo-EM sample preparation

Graphene-oxide (GO) covered grids were prepared following previously published protocols [51], using C-Flat 1.2/1.3 grids (Protochip), and vitrified using a Mark IV Vitrobot (ThermoFisher) at 4°C and 100% humidity. 4 mL of 0.35 mg/mL Csx28 was applied to a freshly prepared GO grid, blotted for 6 seconds and blot force 6, and subsequently plunged into a slurry of liquid ethane cooled using liquid nitrogen.

### Cryo-EM image processing and Model building

Prepared grids were imaged at 63,000X nominal magnification (1.33Å/pixel) using Talos Arctica (ThermoFisher) equipped with a Gatan K3 direct electron detector (Gatan) and Gatan bioquantum energy filter (GIF, Gatan). Prior to image acquisition, the microscope was carefully aligned according to recommended procedures for coma-free and parallel illumination settings [52]. A total of 904 micrographs were collected using SerialEM [53], 2 by 2 image shift, and nominal defocus range of -1 mm to -2.5 mm. The total dose was 47 electrons/Å^2^ fractionated into 44 frames and a total of 2.96 seconds exposure (fig. S6A, Table S1). The following pre-processing and micrograph selection steps were carried out using Warp [54]: beam-induced motion correction, CTF estimation, and particle picking. The final set of 314 micrographs with an estimated resolution of 5 Å or better were imported to cryoSPARC [55] for subsequent processing. Template-based particle picking in cryoSPARC resulted in 247,026 particles. 2D classification of this particle set resulted in multiple 2D classes of side-like views and one top-view with 8- fold symmetry, resulting in 118,793 selected particles. *Ab initio* model generation (C1 symmetry) resulted in a reconstruction with clear 8-fold symmetry. Subsequent downstream processing steps were performed while imposing C8 symmetry (fig. S6B- C).

After non-uniform refinement in cryoSPARC, the particle stack was imported to RELION [56] for subsequent refinement. Further rounds of 3D classification resulted in two major classes of Csx28 octamer, and one minor class consisting of two stacked Csx28 octamers (i.e. a dimer of octamers). As two octamer classes didn’t show distinct conformational differences, two classes of 111,831 particles were combined for further refinement. Extensive CTF refinement and Bayesian polishing with RELION [56, 57] was followed. Subsequent rounds of 3D classification to identify classes with qualitatively high-resolution features improved the final resolution from 4.3 Å to 3.65 Å and 58,694 final particles. Local resolution of the final map was estimated using cryoSPARC (fig. S6D). Although this final map showed significantly improved overall resolution, N-terminal trans-membrane helixes were better resolved in the initial cryoSPARC reconstruction (from 118,793 particles before rounds of classification in RELION, fig. S6C). The appearance of the N-terminal helices (7 Å+ resolution) is likely due to the liberal masking imposed by the initial cryoSPARC reconstruction (which also includes micelle density, 10 Å) and the comparatively worse resolution of the N-terminal helices (∼7 Å) compared to that of the soluble portion (4 – 5 Å) of Csx28 (which was subjected to high-resolution refinement).

The final map was sharpened using RELION postprocess function and automatically estimated B-factor (−118 Å^2^). The sharpened map was of sufficient resolution (3.6 Å) for *de novo* model building (as homologous structures were not available). The initial atomic model was manually built using coot [58], focusing on a single asymmetric unit. Most of the model (from residue 32 to 171) was built using the final map from RELION, and N-terminal trans-membrane helix (from residue 19 to 31) was built using the initial cryoSPARC reconstruction. The manually built subunit was then manually docked into the density using UCSF Chimera [59] in order to create the octameric assembly. This model was subjected to iterative process of manual inspection, followed by manual adjustments using coot [58], followed by symmetric refinement using RosettaES [60]. Model-map FSC (fig. S6E), EMRinger score [61], and MolProbity [62] (Table S1) were calculated using Phenix [63, 64] model validation tools.

### Membrane Fractionation and Western blotting

For western blot detection, Csx28 and Cas13b plasmids were modified to contain a C-terminal V5 and N-terminal 3xHA epitope tag, respectively. Fractionation was performed as previously described [65]. Briefly, 300ml cultures of pEmpty + pACYC184 (ϕλCas13 empty control), HA-Cas13b + pEmpty, pACYC184 + Csx28-V5, and HA-Cas13b + Csx28-V5 were harvested at OD 0.6, resuspended in 15 ml cell resuspension buffer (50 mM sodium phosphate buffer, 300 mM NaCl, 2 mM MgCl_2_, 0.2 mg/ml DNaseI, EDTA-free protease inhibitor (Roche) and 0.1 mg/ml lysozyme) and lysed using a French press at 12,000 psi. Lysate was cleared of unbroken cells by centrifugation (12,000 x g) and samples of the supernatant (whole cell lysate) were collected for Western analysis. The lysate was further purified by ultracentrifugation (180,000 x g) which resulted in cytosol (supernate) and membrane (pellet) fractions. The membrane fraction was resuspended in membrane resuspension buffer (50 mM sodium phosphate buffer, 300 mM NaCl, 5% glycerol, EDTA-free protease inhibitor (Roche), 2% DDM) and was further separated into soluble membrane fraction and insoluble membrane fraction via ultracentrifugation (180,000 x g). Samples from the cytosol, soluble membrane, and insoluble membrane fractions were collected for western analysis. Sample loading was normalized by running 0.1% of each total fraction volume. Two gels with identical sample loading order and amount were probed and stripped for subsequent probing. HA-Cas13b was detected using anti-HA antibody (1:1000 in PBST, Roche) and Csx28- V5 was detected by anti-V5 antibody (1:5000 in PBST, Invitrogen). Cytoplasmic control, DnaK, was detected by rabbit anti-DnaK antibody (1:1000 in PBST, Biorybt) and membrane control, OmpC, was detected by anti-OmpC antibody (1:1000 in PBST, Biorybt).

### *In vivo* Protein crosslinking

Protein *in vivo* crosslinking was carried out with NHS ester, DSS (disuccinimidyl suberate, spacer arm 11.4 Å) (Thermo Scientific) also as suggested by the manufacturer. Briefly, 2×10^10^ non-infected or λ phage 1 hr-post infected bacterial cells expressing V5-tagged Csx28, HA-tagged Cas13 or combination of both proteins together with C3000 host strain as negative control were washed three times in ice-cold 1×PBS (to remove any amine-containing compounds) and treated with 2 mM DSS for 30 min at room temperature. Then reactions were quenched by addition of 1 M Tris pH 7.5 to a final concentration 20 mM for 15 min at room temperature and cells collected by centrifugation. The status of Csx28 protein in crosslinked cells was determined by making the total cell extracts and carrying out Western blot analysis as described above.

### Flow Cytometry

Overnight cultures of *E. coli* C3000 in LB were diluted to OD 0.1 and grown until OD 0.4, at this time cells were infected with λ- phage at an MOI of 1 and 10 mM MgCl_2_ was added to the media. 200 µl samples were collected at time 0 min, 30 min, and 90 min post infection. Samples were treated with the fluorescent voltage sensitive dye DiBAC4(3) (5 µM) for 10 minutes at 37 °C. Treated cells were then spun down and washed with fresh LB+10mM MgCl2 twice and finally diluted 1:10 in fresh LB+10mM MgCl_2_. Samples were measured using a Cytek Aurora Full Spectrum Flow Cytometer measuring 100,000 events (cells) per sample and data were analyzed using FCS Express7 Research (De Novo Software) and data was plotted using GraphPad Prism 9.

### Resazurin assays for cellular viability and metabolic activity

Cells were prepared as previously described as above for bacterial growth curves with the exception that assay was paused after 1 hour post lambda infection to allow for resazurin addition. Resazurin (final concentration 3 µg/ml; stock dissolved in PBS, pH7.4 and then 0.2 µm filtered sterilized) was added to each 200 µl sample-containing well in addition to three wells of LB media control, the assay was resumed with fluorescent measurements taken every 10 minutes with 560 nm excitation / 590 nm emission wavelengths using a Tecan Spark plater reader.

## Acknowledgments

We thank K. Awayda, J. Nicosia, and N.Tong for insightful advice on experiments conducted for this manuscript, and the O’Connell and Kellogg labs for helpful discussions. We thank M. Dumont for invaluable advice on membrane protein purification and access to light scattering measurements. We thank C. Cavender for assistance with statistical analyses. We thank M. Cochran and the URMC Flow Cytometry Resource for assistance with the flow cytometry experiments. We thank the Cornell Center for Materials Research facility, as well as K. Spoth and M. Silvestry-Ramos, for maintenance of electron microscopes used for this research (NSF MRSEC program, DMR-1719875); XSEDE for computational resources used for image processing (MCB200090 to E.H.K.). We thank Sam Sternberg for critical feedback on the manuscript.

## Author contributions

M.R.O and A.R.V conceived the project. A.R.V carried out the phage assays, A.R.V carried out protein purification. M.R.O carried out the SEC-SLS experiments. J.-U.P. prepared samples for cryo-EM imaging; J.-U.P. analyzed, processed, and refined cryo-EM images to obtain 3D reconstructions; J.-U.P., and E.H.K. built and refined atomic models. A.R.V and B.P carried out the membrane fractionation and *in vivo* crosslinking experiments. A.R.V. carried out the flow cytometry experiments with assistance from M.R.O. A.R.V carried out the resazurin experiments with assistance from M.R.O. M.R.O and A.R.V drafted the manuscript with input from J.- UP, E.H.K and B.P, and all authors synthesized the ideas, contributed to figures, and reviewed and edited the final manuscript.

## Data and materials availability

Atomic models are available through the Protein Data Bank (PDB) with accession codes 7S92 (Csx28); cryo-EM reconstructions are available through the EMDB with accession codes EMD-24929 (Csx28).

## Supplementary Materials (see separate pdf)

Figs. S1 to S11

Tables S1 to S3

