## Supplementary Material for "CRISPR-Csx28 forms a Cas13b-activated membrane pore required for robust CRISPR-Cas adaptive immunity"

**This PDF file includes:**

Figs. S1 to S11
Tables S1 to S2

**Supplementary Figures**

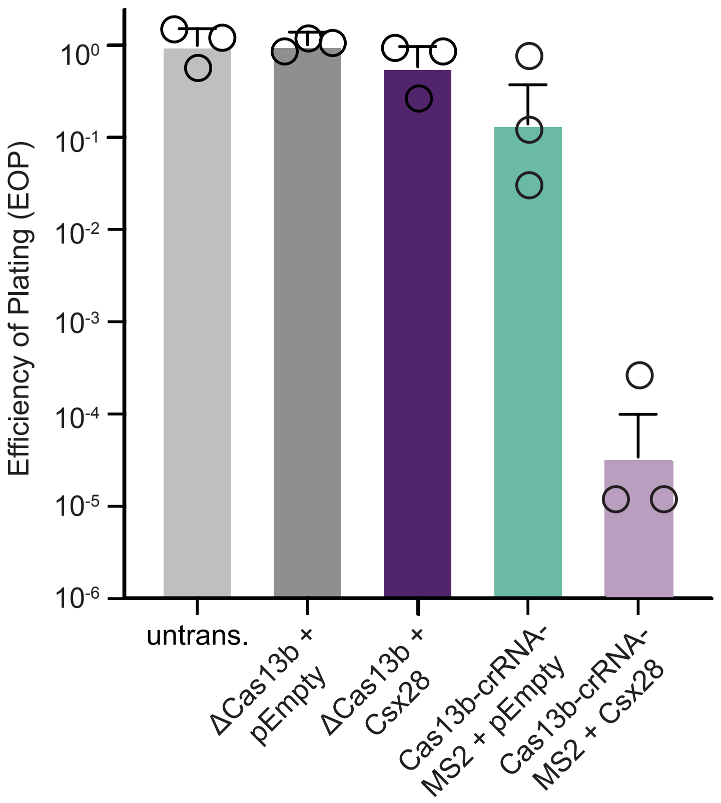

**Fig. S1. Csx28 significantly enhances Type VI-B2 CRISPR-Cas immunity against MS2 ssRNA phage and requires targeting Cas13b for this activity.** Efficiency of Plating (EOP) assays measuring MS2-phage infection susceptibility of untransformed (untrans.) *E. coli* or strains carrying the indicated Cas13b-crRNA-MS2, Csx28, or pEmpty plasmids. ΔCas13b is no Cas13b control. Data is shown as mean ± s.e.m for $n = 3$ biological replicates.

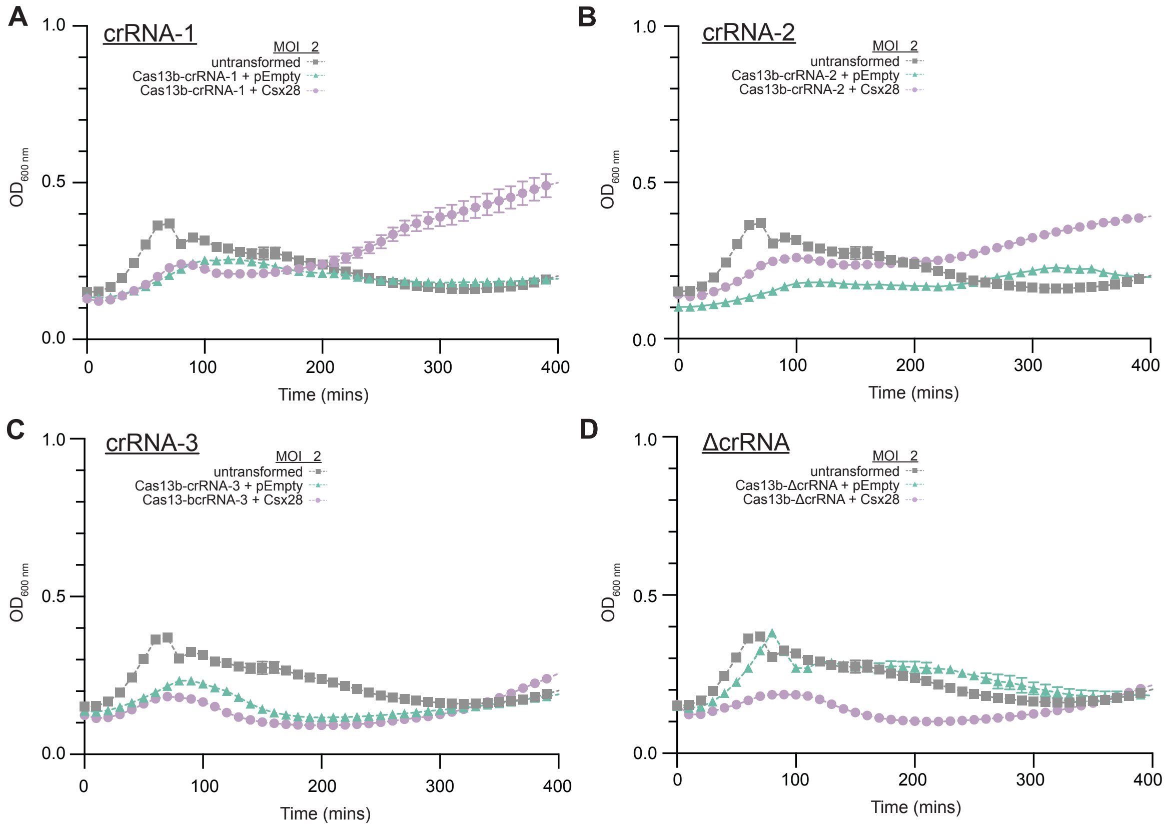

**Fig. S2. Csx28 also allows *E. coli* to recover from λ-phage infection at high MOIs.** Growth curves of *E. coli* strains carrying the indicated Cas13b-crRNA-X, Csx28, or pEmpty plasmids, as measured using $OD_{600}$ after the addition of λ-phage at an MOI of 2. (**A-D**) are growth curves for crRNA-1, crRNA-2, crRNA-3 and ΔcrRNA- containing strains, respectively. Data is shown as mean ± s.e.m for *n* = 3 biological replicates.

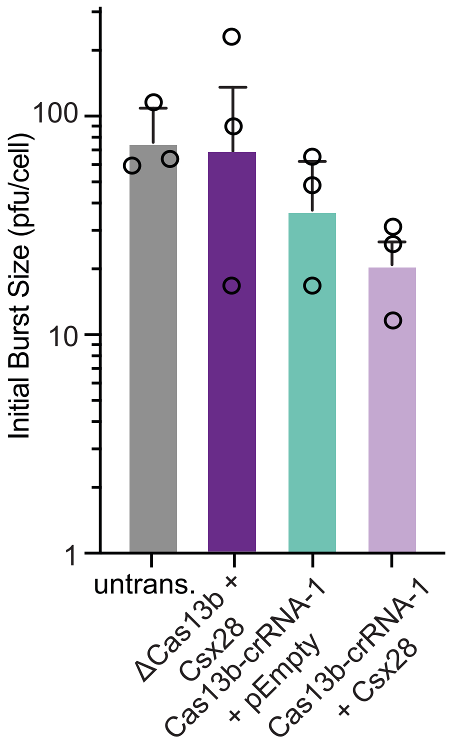

**Fig. S3. Cas13b-activated Csx28 reduces initial phage burst size.** One-step phage growth (burst) assays measuring initial λ-phage production from one round of phage replication for untransformed (untrans.) *E. coli* strains carrying the indicated Cas13b-crRNA-1, Csx28-containing, or pEmpty plasmids infected with λ-phage at an MOI of 0.1. Data is shown as mean ± s.e.m for $n$ = 3 biological replicates.

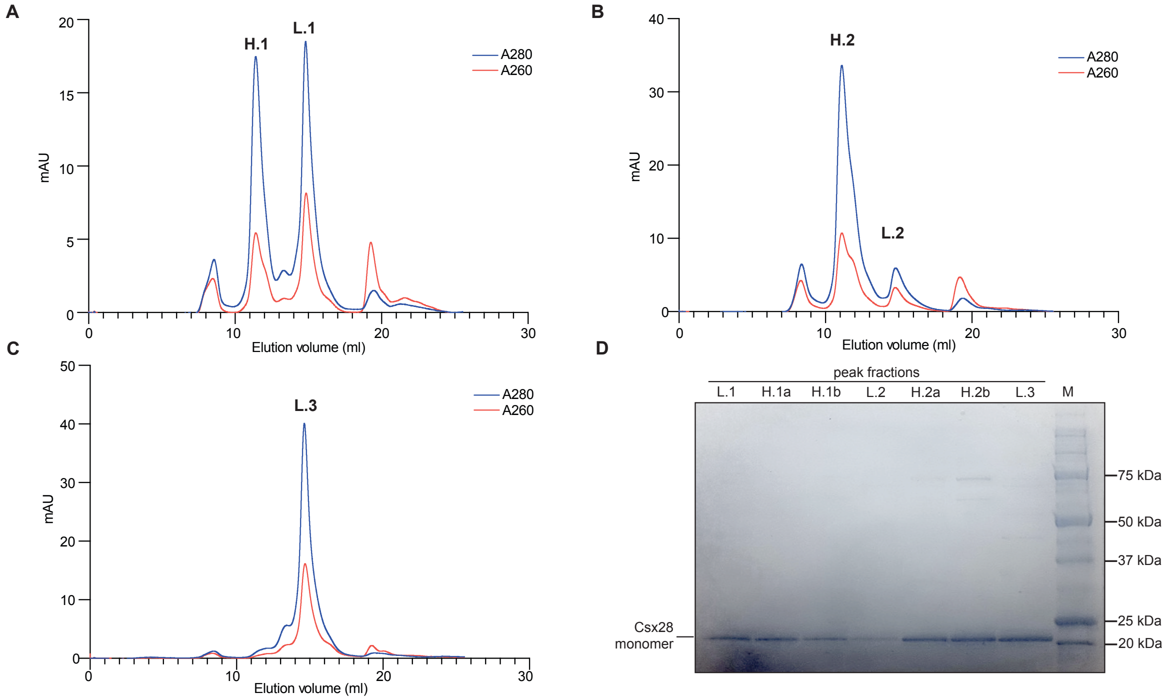

**Fig S4. Full length Csx28 can be purified from *E. coli* to homogeneity and exists primarily in two distinct non-exchanging oligomeric states.** (**A**) Size exclusion chromatography trace of Csx28. "Light" and "Heavy" peaks are labeled as L.1 and H.1, respectively. These peaks were separately pooled and concentrated, and then subjected to an additional size exclusion cleanup step to generate (**B**) for the Heavy fraction and (**C**) for the Light fraction. The Heavy and residual Light peak in (B) are labeled "H.2" and "L.2", respectively and in (C) the Light peak is labeled L.3. These were all separately pooled and concentrated for subsequent experiments. (**D**) SDS-PAGE analysis of fractions from (A-C). H.1a, H.1b, and H.2a and H.2b were sampled from two separate fractions in the H.1 peak in (A) and H.2 peak in (B), respectively, prior to pooling the fractions.

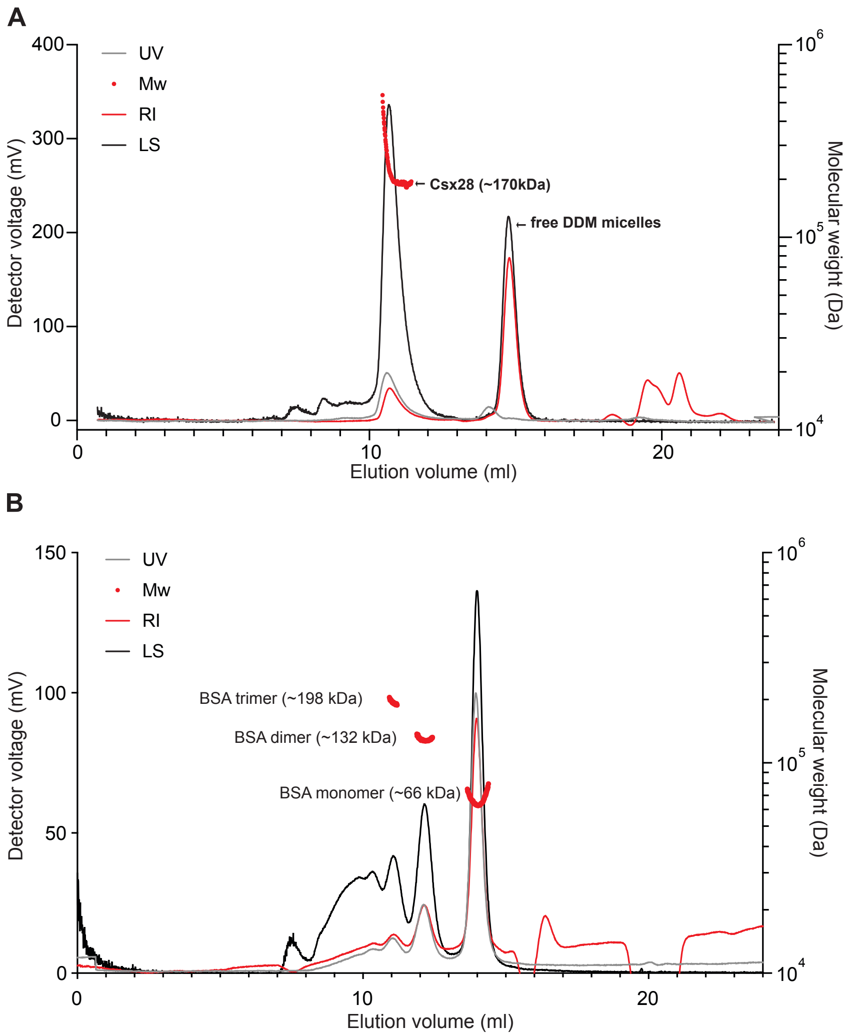

**Fig. S5. SLS-SEC reveals that Csx28 forms octamers in solution that are in intermediate exchange with larger oligomers over the course of the SEC run.** Three detector (UV: UV 280 nm absorbance, RI: refractive index, LS: static light scattering) traces for (**A**) Csx28 and (**B**) BSA, as a SLS-SEC sizing control and reference. Molecular weights were determined using the three-detector method (*48*). The experimentally determined molecular weights of each peak are shown.

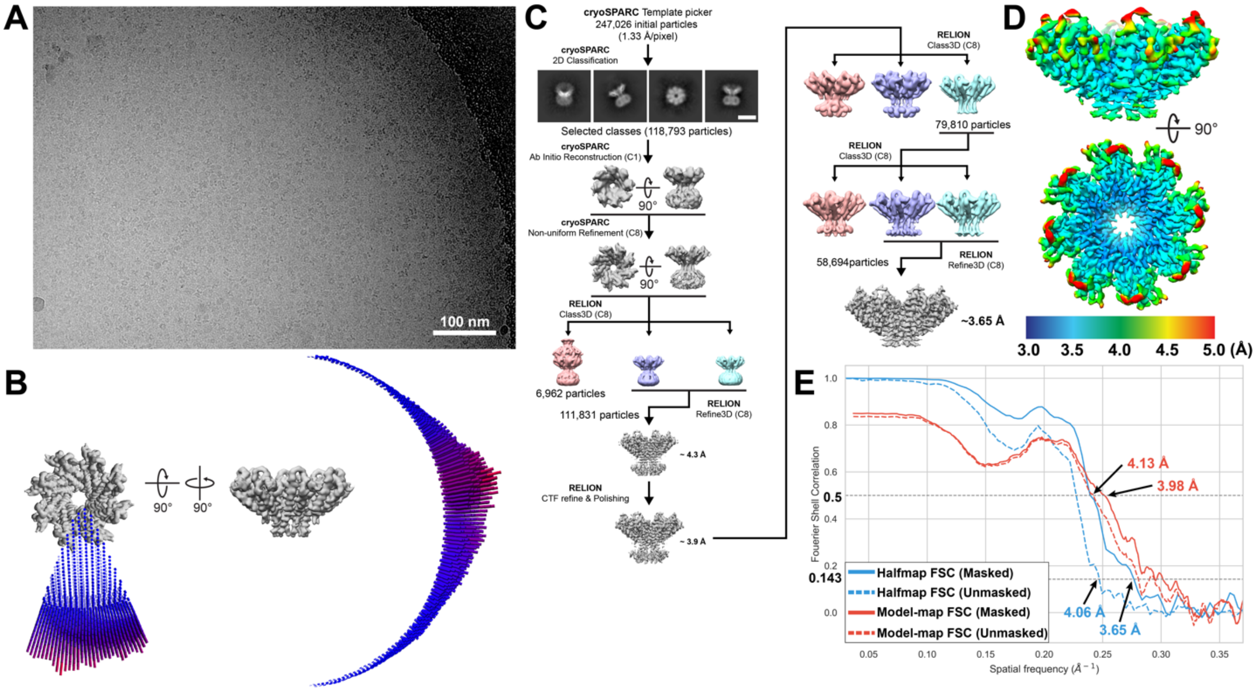

**Fig. S6. Cryo-EM data collection and image processing of a Csx28 octameric structure.**

(**A**) Representative micrograph from graphene oxide covered cryo-EM grid of Csx28. Images were taken using Talos Arctica equipped with a K3 direct electron detector. Scale bar represents 100 nm. (**B**)**.** Euler angle orientation distribution of the final reconstruction of Csx28 using C8 symmetry in RELION (*49*). (**C**) Image processing workflow for the Csx28 octamer. Scale bar in 2D class average represents 100 Å. Since 2D average of the top-view, and the initial *ab initio* reconstruction showed clear 8-fold rotational symmetry, C8 symmetry was applied in the following refinements. (**D**) Local resolution of Csx28 octamer is estimated using cryoSPARC (*50*). Legend on the bottom of the panel represents the corresponding resolution for the surface color of the reconstruction. (**E**) Fourier shell correlation (FSC) plot of the Csx28 reconstruction and model. Solid or dashed lines indicate masked or unmasked FSC curves, respectively. FSC between half-maps (blue) or between full-map and atomic model (red) is presented with the estimated resolution using the corresponding cutoff (0.143 for half-maps, 0.5 for model-map FSC).

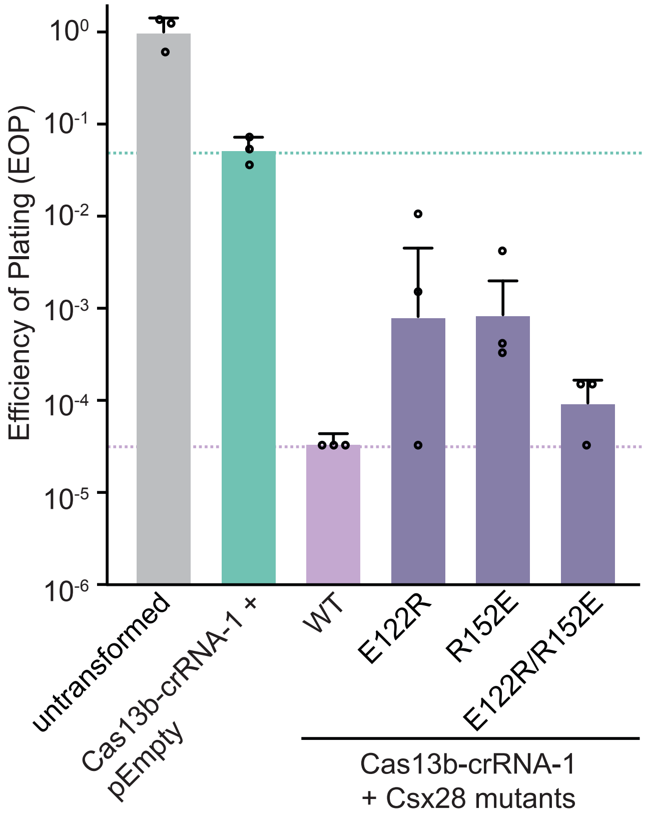

**Fig. S7. Charge swap analysis of the potential E122:R152 salt bridge provides further evidence that R152 is engaged in a salt bridge and likely not playing a role in the Csx28 protomer:promoter interface.** Efficiency of Plating (EOP) assays measuring λ-phage infection susceptibility of *E. coli* strains carrying the indicated Cas13b-crRNA-1, and WT or mutant Csx28 plasmids. Data is shown as mean ± s.e.m for $n$ = 3 biological replicates.

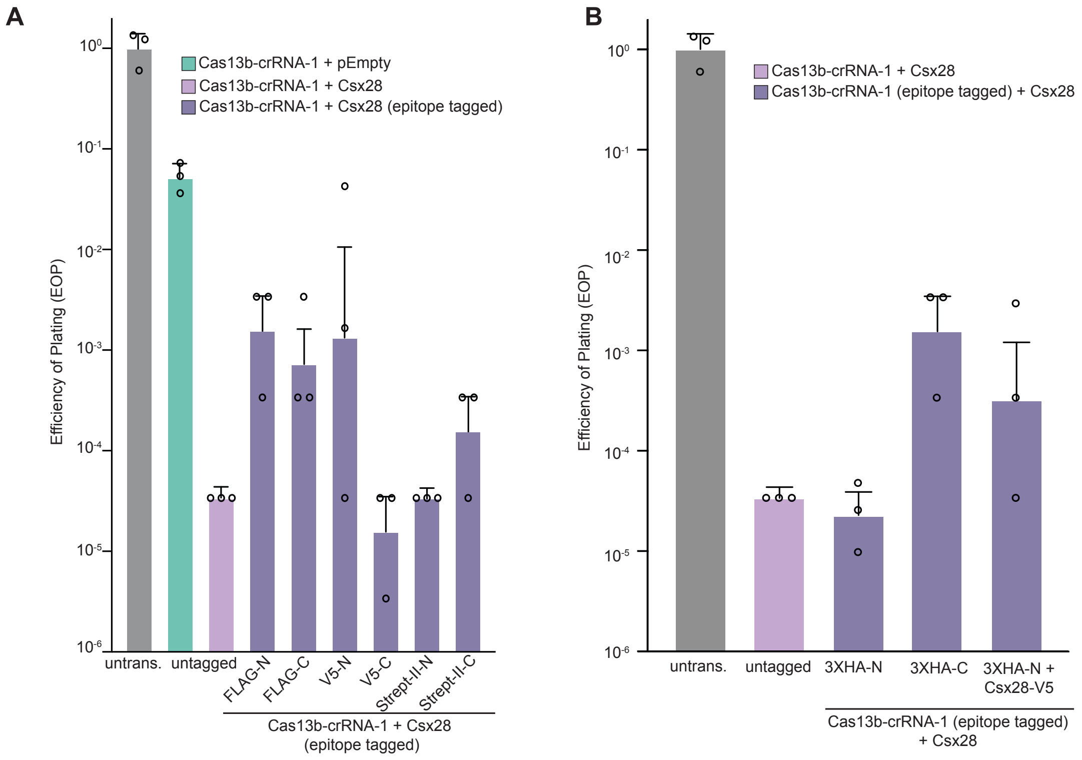

**Fig. S8. Cas13b and Csx28 anti-phage defense activity is differentially affected by the choice of epitope tag sequence and location.** (**A**) Efficiency of Plating (EOP) assays measuring λ-phage infection susceptibility of untransformed (untrans.) *E. coli* or strains carrying the indicated Cas13b-crRNA-1 and WT or epitope-tagged Csx28 plasmids. (**B**) Efficiency of Plating (EOP) assays measuring λ-phage infection susceptibility of *E. coli* strains carrying the indicated epitope-tagged Cas13b-crRNA-1 and epitope tagged Csx28 plasmids. Data is shown as mean ± s.e.m for *n* = 3 biological replicates.

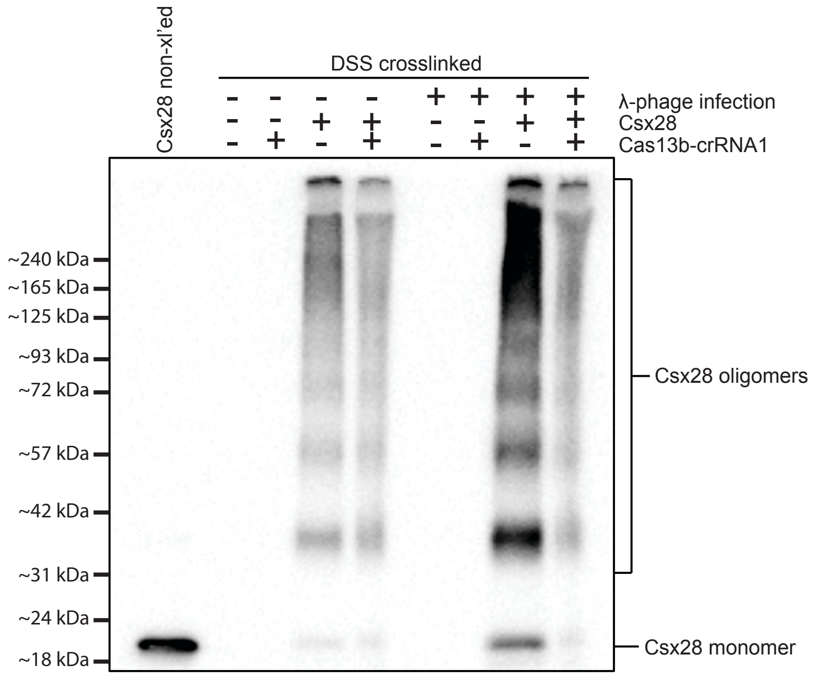

**Fig. S9. Csx28 can form oligomers *in vivo*.** A Western blot to detect Csx28 crosslinking *in vivo*. Cas13b-crRNA-1 and V5-tagged Csx28 expressing *E coli* were subject to DSS crosslinking in culture, were subsequently lysed and total cell lysates were subjected to SDS-PAGE prior to Western blot analyzes. A non-crosslinked Csx28-expressing total cell lysate (Csx28 non-xl'ed) was used as a control. Blots were probed with an anti-V5 antibody to detect Csx28-V5.

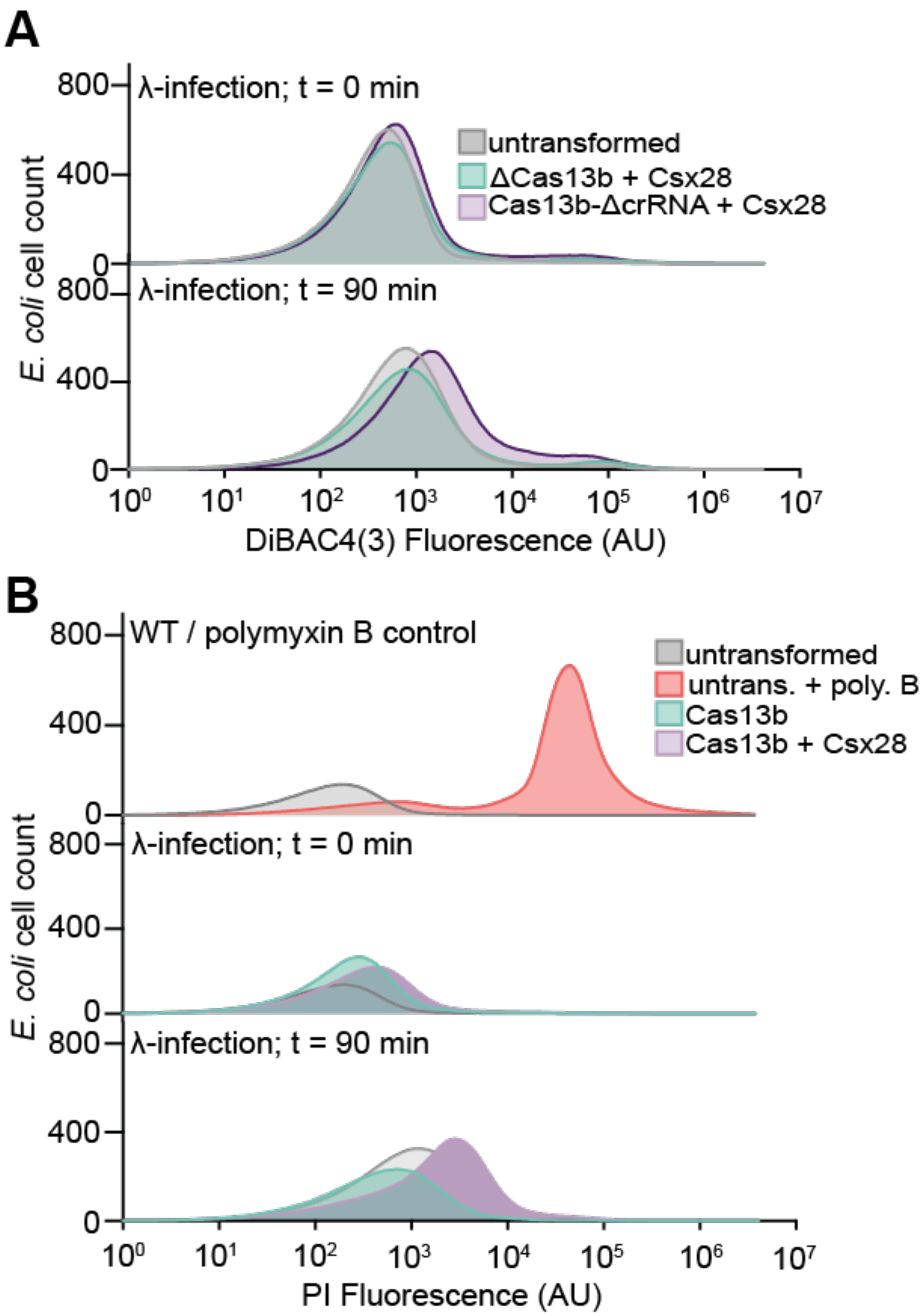

**Fig. S10. Csx28 expression alone cannot induce membrane depolarization and Csx28 activation doesn't lead to a substantial uptake of propidium iodide.** (**A**) Flow cytometry histograms of a DiBAC4(3) staining assay measuring membrane depolarization of untransformed *E. coli or E. coli* strains expressing Cas13b-crRNA-1 and Csx28, or Cas13b-ΔcrRNA and Csx28 over the course of a λ-phage infection (MOI of 1). (**B**) Flow cytometry histograms of a propidium iodide (PI) staining assay measuring gross membrane integrity of untransformed (untrans.) *E. coli or E. coli* strains expressing Cas13b-crRNA-1, or Cas13b-crRNA-1 and Csx28 over the course of a λ-phage infection (MOI of 1). A Polymyxin B (Poly. B; a membrane integrity disrupter)

treated *E. coli* sample was used as positive control for loss of membrane integrity. In all samples above, 10,000 *E. coli* cells were analyzed.

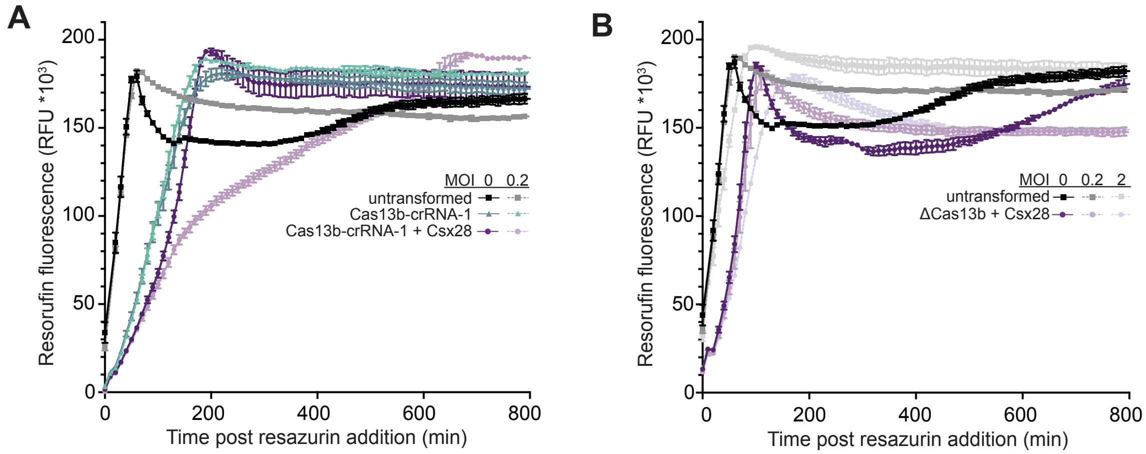

**Fig. S11. Activated Csx28 results in loss of metabolic activity at low MOI infection and requires Cas13b-mediated activation for this effect.** Resazurin metabolism assay measuring the conversion of resazurin to fluorescent resorufin over time as a function of active respiration for (**A**) untransformed *E. coli or E. coli* strains expressing Cas13b-crRNA-1, or Cas13b-crRNA-1 and Csx28 either in the absence or presence of λ-phage infection (MOI of 0.2), or (**B**) untransformed *E. coli or E. coli* strains expressing Cas13b-crRNA-1 and Csx28 either in in the absence or presence of λ-phage infection (MOI of 0.2 and 2). Resazurin is added to the growing cultures at one hour post λ-phage infection. Data is shown as mean ± s.e.m for $n$ = 3 biological replicates.

**Table S1. Cryo-EM Statistics for Data Collection and Model Refinement.**

| | |
|---|---|
| Name | Csx28 octamer |
| PDB ID | 7S92 |
| EMDB ID | EMD-24929 |
| **Data collection and Processing** | |
| Microscope | Talos-Arctica |
| Voltage (keV) | 200 |
| Camera | K3 |
| Magnification | 63,000 |
| Pixel size at detector (Å/pixel) | 1.33 |
| Total electron exposure ($e^{-}$/Å$^2$) | 47 |
| Exposure rate (e-/pixel/sec) | 28.09 |
| Number of frames collected during exposure | 44 |
| Defocus range (mm) | -1 – -2.5 |
| Automation software | SerialEM |
| Energy filter slit width | 20 keV |
| Micrographs collected (no.) | 904 |
| Micrographs used (no.) | 314 |
| Total extracted particles (no.) | 247,026 |
| **For each reconstruction:** | |
| Refined particles (no.) | 118,793 |
| Final particles (no.) | 58,694 |
| Symmetry | C8 |
| Resolution (global, Å) | |
| FSC 0.5 (unmasked/masked) | 4.43/4.18 |
| FSC 0.143 (unmasked/masked) | 4.06/3.65 |
| Resolution range (local, Å) | 3.5 – 5 |
| Resolution range due to anisotropy (Å) | 3.5 – 3.8 |
| Map sharpening *B* factor (Å$^2$) | -118 |
| Map sharpening methods | RELION |
| **Model composition** | |
| Protein residues | 1,152 |
| Ligands | 0 |
| RNA/DNA | 0 |
| **Model Refinement** | |
| Refinement package | Coot/Rosetta |
| - real or reciprocal space | Real |
| - resolution cutoff (Å) | 3.5 |

| | |
|---|---|
| Model-Map scores | |
| - CC | 0.79 |
| - Average FSC (0.5 cutoff, Å) | 3.98 |
| R.m.s deviations from ideal values | |
| Bond length (Å) | 0.017 |
| Bond angles (°) | 1.483 |
| **Validation** | |
| MolProbity score | 1.80 |
| CaBLAM outliers (%) | 4.07 |
| Clashscore | 9.89 |
| Poor rotamers (%) | 0 |
| C-beta outliers (%) | 1.06 |
| EMRinger score | 2.14 |
| Ramachandran plot | |
| Favored (%) | 95.9 |
| Outliers (%) | 0.0 |

**Table S2. Plasmids used in this study.**

| Plasmid | Description | Source |
|---|---|---|
| pPbuCas13b | Lac inducible promoter adaptation of addgene #89906 (cam) | This study |
| pPbuCas13b-ΔcrRNA | Deletion of crRNA and promoter region | This study |
| pPbuCas13b-crRNA1 | Spacer targeting λ N gene | This study |
| pPbuCas13b$^{dHEPN}$-crRNA1 | Deactivated nuclease activity of HEPN | This study |
| pPbuCas13b-crRNA2 | Spacer targeting λ Q gene | This study |
| pPbuCas13b-crRNA3 | Spacer targeting λ R gene | This study |
| pPbuCas13b-crRNA-MS2 | Spacer targeting MS2 replicase gene | This study |
| pΔPbuCas13b | Deletion of Cas13b orf | This study |
| pHA-PbuCas13b-crRNA1 | N-terminal HA tag | This study |
| pPbuCas13b-crRNA1-HA | C-terminal HA tag | This study |
| pPbuCsx28 | Lac inducible promoter adaptation of addgene #89909 (amp) | This study |
| pEmpty | Deletion of Csx28 orf and promoter region | This study |
| pPbuCsx28-Y55A | Conserved structure-guided guided mutant Y55A | This study |
| pPbuCsx28-T62A | Conserved structure-guided guided mutant T62A | This study |
| pPbuCsx28-Y104A | Conserved structure-guided guided mutant Y104A | This study |
| pPbuCsx28-Y104F | Conserved structure-guided guided mutant Y104F | This study |
| pPbuCsx28-E122R | Structure-guided guided mutant E122R | This study |
| pPbuCsx28-R152A | Predicted HEPN residue mutant R152A | This study |
| pPbuCsx28-R152E | Structure-guided guided mutant R152E | This study |
| pPbuCsx28-H157A | Predicted HEPN residue mutant H157A | This study |
| pPbuCsx28-R165A | Structure-guided guided mutant R165A | This study |
| pPbuCsx28-R165E | Structure-guided guided mutant R165E | This study |
| pPbuCsx28-Y104A/H157A | Y104A & H157A double mutant | This study |
| pPbuCsx28-Y104A/R165A | Y104A & R165A double mutant | This study |
| pPbuCsx28-R152A/H157A | R152A & H157A double mutant | This study |

| | | |
|---|---|---|
| pPbuCsx28-H157A/R165A | H157A & R165A double mutant | This study |
| pFlag-PbuCsx28 | N-terminal Flag tag | This study |
| pPbuCsx28-Flag | C-terminal Flag tag | This study |
| pStrepII-PbuCsx28 | N-terminal Strep II tag | This study |
| pPbuCsx28-StrepII | C-terminal Strep II tag | This study |
| pV5-PbuCsx28 | N-terminal V5 tag | This study |
| pPbuCsx28-V5 | C-terminal V5 tag | This study |
| p2CcT-PbuCsx28 | PbuCsx28-MBP overexpression | This study |

**Table S3. Oligonucleotide primers/gene fragments used in this study.**

| Primer | Sequence |
|---|---|
| PbuCsx28_Lac Fix_F | atggattatatggaattagcgaaagaggcat |
| PbuCsx28_Lac Fix_R | agctgtttcctgtgtgaaattgttatcc |
| Lac sequence_F | acgatataagttgtaattctcatgtgcaattaatgtgagttagctcactcattag g |
| Lac sequence_R | caaataatttatcttgtttttgcatagctgtttcctgtgtgaaattgt |
| PbuCas13b_Lac Fix_F | atgcaaaaacaagataaattatttgtagatagaaagaaaaatgct |
| PbuCas13b_Lac Fix_R | acatgagaattacaacttatatcgtatggggc |
| λ crRNA-1_F | acaacagctaatccggaatcgcacttacggccaa |
| λ crRNA-1_R | caacttggccgtaagtgcgattccggattagctg |
| λ crRNA-2_F | acaaggaaagagagtcagaagccgtggcccgtgg |
| λ crRNA-2_R | caacccacgggccacggcttctgactctctttcc |
| λ crRNA-3_F | acaaagatattgctgcaacggtcgattgcctgac |
| λ crRNA-3_R | caacgtcaggcaatcgaccgttgcagcaatatct |
| MS2 crRNA-1_F | acaacgagagaaagatcgcgaggaagatcaatac |
| MS2 crRNA-1_R | caacgtattgatcttcctcgcgatctttctctcg |
| PbuCsx28_ ΔcrRNA _F | atcgtccattccgacagc |
| PbuCsx28_ ΔcrRNA _R | ttagttttattttctgatttttcaatttcttttatggc |
| ΔPbuCas13b_F | attgacagctagctcagtccta |
| ΔPbuCas13b_R | acatgagaattacaacttatatcgtatggggc |
| dPbuCas13b_F1 | tattctgaagaatccccaaaaccgatatttgaaacaagcc |
| dPbuCas13b_R1 | cttgtacgcggagtagtagttcgcgagaacctgaag |
| dPbuCas13b_F2 | tatcccatgtatgatgccacactctttgctgaagt |
| dPbuCas13b_R2 | ctggttcgcggaaaaagcattcgccactgcaatc |
| ΔPbuCsx28_F | ccgggtaccgagctcgaattc |
| ΔPbuCsx28_R | gcgttgcgctcactgccc |
| PbuCsx28_Y55A_F | aagacagcatattgatttgggaaaaaacaaaagatggtaagc |
| PbuCsx28_Y55A_R | cattttcagtatttgtagtaaatctgagtagtttcgccaggc |
| PbuCsx28_T62A_F | aagacagcatattgatttgggaaaaaacaaaagatggtaagc |
| PbuCsx28_T62A_R | cattttcagtattcgcagtaaatctgagtagtttatacaggc |
| PbuCsx28_Y104A_F | gatgtggtatttttatgtcaaaagaggctctgtcactaatttcc |
| PbuCsx28_Y104A_R | cttcgttcgctatttctttggaaagctcttttataaatttgcg |
| PbuCsx28_Y104F_F | gatgtggtatttttatgtcaaaagaggctctgtcactaatttcc |
| PbuCsx28_Y104F_R | cttcgttaaatatttctttggaaagctcttttataaatttgcg |
| PbuCsx28_E122R_F | gtttcatgctttccgcccaaaacaatcc |
| PbuCsx28_E122R_R | cataaacaatatttctatagcgggaaattagtgacagagcctc |
| PbuCsx28_R152A_F | tcgatagaaatcagacaagcaatcaatctgaaaaaacgag |
| PbuCsx28_R152A_R | gaggttttggtgaatctttttcatcgcttctacagactc |
| PbuCsx28_R152E_F | tcgatagaaatcagacaagcaatcaatctgaaaaaacgag |
| PbuCsx28_R152E_R | gaggttttggtgaatctttttcatttcttctacagactc |
| PbuCsx28_H157A_F | tcgatagaaatcagacaagcaatcaatctgaaaaaacgag |
| PbuCsx28_H157A_R | gaggttttgcgcaatctttttcattcgttctacagactc |
| PbuCsx28_R165A_F | aatctgaaaaaacgagatttaagattttaaccgggtaccg |

| | |
|---|---|
| PbuCsx28_R165A_R | gattgcttgcgcgatttctatcgagaggttttgg |
| PbuCsx28_R165E_F | aatctgaaaaaacgagatttaagattttaaccgggtaccg |
| PbuCsx28_R165E_R | gattgcttgttcgatttctatcgagaggttttgg |
| 1x_Flag_PbuCsx28_F | acaaaggctcttctggttcggattatatggaattagcgaaagaggc |
| 1x_Flag_PbuCsx28_R | cgtcgtcgtccttgtagtccatagctgtttcctgtgtgaaattg |
| PbuCsx28_1x_Flag_F | caaggacgacgacgacaaataaccgggtaccgagct |
| PbuCsx28_1x_Flag_R | tagtccgaaccagaagagccaaatcttaaatctcgtttttcagattg |
| StrepII_PbuCsx28_F | ggtgggggctcttctggttcggattatatggaattagcgaaagaggc |
| StrepII_PbuCsx28_R | cggttgaggatgtgcccacgccatagctgtttcctgtgtgaaattg |
| PbuCsx28_StrepII_F | gcacatcctcaaccgggtgggtaaccgggtaccgagct |
| PbuCsx28_StrepII_R | ccacgccgaaccagaagagccaaatcttaaatctcgtttttcagattg |
| V5_PbuCsx28_F | gtctcgattctacgggctcttctggttcggattatatggaattagcgaaagagg c |
| V5_PbuCsx28_R | cgaggagagggttagggataggcttacccatagctgtttcctgtgtgaaattg |
| PbuCsx28_V5_F | taaccctctcctcggtctcgattctacgtaaccgggtaccgagct |
| PbuCsx28_V5_R | gggataggcttacccgaaccagaagagccaaatcttaaatctcgtttttcag attg |
| 3x_HA_PbuCas13b_F | cgcatacccgtatgacgttccggactatgccgggtcaagtggctctcaaaa acaagataaattatttgtagatagaaagaaaaatg |
| 3x_HA_PbuCas13b_R | tagtctggaacgtcgtatggataagcgtagtcagggacatcataagggtac atagctgtttcctgtgtgaaattg |
| PbuCas13b_3x_HA_F | cgacgttccagactacgcatacccgtatgacgttccggactatgcctaaatt gacagctagctcagtcc |
| PbuCas13b_3x_HA_R | tatggataagcgtagtcagggacatcataagggtaagagccacttgaccc gtttttattttctgatttttcaatttcttttatggc |
